# Small non-coding RNA content in plasma-derived extracellular vesicles distinguish ataxic SCA3 mutation carriers from pre-ataxic and control subjects

**DOI:** 10.1101/2024.01.04.574044

**Authors:** Magda M Santana, Patrick Silva, Maria M Pinto, Laetitia Gaspar, Rui Nobre, Sónia Duarte, Tânia Monteiro Marques, Margarida Gama-Carvalho, Cristina Januário, Inês Cunha, Joana Afonso Ribeiro, Jeannette Hübener-Schmid, Jon Infante, Mafalda Raposo, Manuela Lima, Hector Garcia-Moreno, Paola Giunti, Bart van de Warrenburg, Matthis Synofzik, Jennifer Faber, Thomas Klockgether, ESMI Study Group, Luís Pereira de Almeida

## Abstract

Spinocerebellar ataxia type 3 (SCA3), a neurodegenerative disorder caused by a CAG expansion in the *ATXN3* gene, is the most common spinocerebellar ataxia subtype worldwide. Currently, there is no therapy to stop or prevent disease progression. Promising therapeutic strategies are emerging, but their translation into clinical practice requires sensitive and reliable biomarkers. Blood circulating extracellular vesicles constitute a promising source of biomarkers with potential to track alterations of the central nervous system due to their ability to cross the blood brain barrier.

Here, we perform sequencing analysis of small RNAs from plasma-derived extracellular vesicles from SCA3 mutation carriers (10 pre-ataxic and 10 ataxic) and 12 control subjects to identify potential RNA biomarker candidates for this disease.

Data showed that plasma-derived extracellular vesicles from ataxic SCA3 mutation carriers are enriched in mitochondrial, nuclear, and nucleolar RNA biotypes compared to pre-ataxic and control subjects. Moreover, ataxic mutation carriers could be discriminated from control and pre-ataxic subjects based on the miRNAs or piRNAs content, but not tRNA. Furthermore, we identified a subset of differentially expressed miRNAs and piRNAs that clearly differentiate ataxic mutation carriers from pre-ataxic and control subjects.

These findings open new avenues for further investigation on the role of these RNAs in the pathogenesis of SCA3 and their potential as biomarkers for this disease.

## Introduction

Spinocerebellar ataxias (SCAs) are a group of more than 40 subtypes of autosomal dominantly inherited neurodegenerative diseases [1]. SCAs are commonly characterized by a progressive loss of balance and coordination resultant from the degeneration of specific regions of the central nervous system, such as the cerebellum, brainstem, basal ganglia and spinal cord [2]. Spinocerebellar ataxia type 3 (SCA3), also referred to as Machado-Joseph disease (MJD), is the most predominant SCA subtype worldwide [3]. SCA3 is caused by a pathological expansion of the cytosine-adenosine-guanine (CAG) trinucleotide repeat in the coding region of the *ATXN3* gene, located at the 14q12.32 chromosomal locus [4]. SCA3 symptoms are usually manifested during adulthood – age of symptoms onset is highly variable but most commonly between the second and fifth decade of life [9] –, and the disease progresses relentlessly to the point that patients become wheelchair-bound and need assistance to perform daily tasks [10]. Although the past decade has seen a vast advancement in the development of novel therapies for this and other polyQ disorders [11, 12], an effective treatment that counteracts SCA3 or relieves its progression is still lacking. Most therapeutic strategies have failed to provide meaningful results in clinical trials partly due to the use of subjective clinical scales that lack sensitivity for monitoring disease fluctuations. Therefore, there is a pressing need to identify objective and sensitive biomarkers for SCA3 that would facilitate the monitoring of disease progression and the assessment of therapy response in forthcoming interventional studies. This has been one of the main focuses of research in SCA3, with initiatives such as the European Spinocerebellar Ataxia Type 3/Machado-Joseph Disease Initiative (ESMI) contributing to substantial progress in this area [13–17].

Extracellular vesicles (EVs) are nanosized (30-1000 nm in diameter), membrane-surrounded, structures shed by nearly all cell types into the extracellular environment [18]. These structures have been shown to carry a panoply of molecules that reflect the biomolecular composition of tissues/cells of origin, including proteins, lipids, and RNA species [18, 19]. EVs are very attractive for biomarker research because (1) they can cross the blood-brain barrier, allowing brain-released EVs to be found in easily accessed peripheral samples [20]; (2) their cargo is protected from degradation by the lipid bilayer, remaining stable in circulation [21]; and (3) they may harbor unique signatures of pathological states, providing evidence of disease and disease progression [22]. One of the most widely studied molecules carried by EVs are RNAs, particularly, small RNAs such as microRNAs. These are particularly attractive biomarkers given their involvement in multiple biological processes in both physiological and disease conditions. Indeed, several studies have reported alterations in EV-RNAs in the context of various neurodegenerative disorders, such as Parkinson’s disease [23] and amyotrophic lateral sclerosis [24], reinforcing the potential of EV-RNAs as disease biomarkers.

In this study, we isolated circulating EVs from plasma samples obtained from molecularly confirmed SCA3 pre-ataxic and ataxic subjects, as well as healthy controls, and performed small RNA-sequencing (sRNA-seq) to identify potential RNA biomarker candidates for SCA3.

## Materials and methods

### Ethical approval

This study was conducted by the European Spinocerebellar ataxia type-3/Machado-Joseph Disease Initiative (ESMI) upon the approval of local ethical committees of all participating centers. Informed consent was obtained from all participants prior to enrolment, in accordance with the Declaration of Helsinki.

### Study design

Molecularly confirmed SCA3 mutation carriers (*ATXN3* repeat length ≥ 60 [25]) and control subjects (molecularly excluded for the *ATXN3* expansion, without neurological disease) were recruited by the ESMI partners at the respective research centers. Demographic information and clinical data were collected from all participants, including age, sex and the score for the Scale for the Assessment and Rating of Ataxia (SARA) [26]. *ATXN3* genotype was determined by PCR-based fragment length analysis, in a centralized manner, for all participants. SCA3 mutation carriers were classified according to SARA score as pre-ataxic (SARA score < 3) or ataxic (SARA score ≥ 3). Individuals without neurological disease were included as control subjects. An effort was made to ensure a balanced distribution of age and gender and minimize pre-analytical and biological biases in the three biological groups. The ataxic group was selected to portray mid-stage SCA3, characterized by moderate ataxia, to make sure significant differences were observed compared to the pre-ataxic stage.

### Blood collection and processing

Peripheral blood was collected from each participant according to a standardized protocol implemented at all participating research centers [27]. Briefly, approximately 8.5 mL of blood was drawn into a plasma preparation tube (362799, BD Vacutainer) using a 21G butterfly needle (BG2134, MediPlus). Tubes were mixed by inversion (8-10 times) and centrifuged at 1000 x g for 10 min at room temperature for plasma separation. Plasma samples were then filtered with a 0.8 µm filter (SLAA033SB, Merck Millipore), aliquoted into a microcentrifuge tube, and stored at -80 °C until further analysis.

### EV isolation from plasma

EVs were isolated from plasma by size exclusion chromatography (SEC), following a protocol previously optimized [28]. In short, 1000 µL of filtered plasma samples were thawed at 37 °C and centrifuged at 10.000 x g for 10 minutes. The supernatant was collected and 900 µL were loaded into qEV columns (#SP1, Izon Science). Fractions (F) of 500 µL were collected with an Automatic Fraction Collector (AFC, Izon Science), and EV-rich fractions (F8-F10, as previously optimized in [28]) were pooled (1500 µL total) and stored at -80°C until analysis.

### Evaluation of particle size and concentration by nanoparticle tracking analysis (NTA)

The size and concentration of isolated plasma-derived EVs were measured using a NanoSight NS300 (Malvern Panalytical, Malvern, the United Kingdom) with a 488 nm laser and a sCMOS camera module. The instrument was operated according to the manufacturer’s instructions. Five 60-second videos were recorded from each sample using a camera setting of 13-14. Data was analyzed using NTA 3.2 software (Malvern Panalytical) with a detection threshold of 3.

### RNA extraction and small RNA sequencing

Total RNA was extracted from plasma-derived EVs (F8-F10, 1500 µL total) using the Exosomal RNA Isolation Kit (#58000, Norgen Biotek Corp.) The purified RNA was concentrated using Norgen’s RNA Clean-Up and Concentration Kit (Cat. 23600) and quantified using the Agilent RNA 6000 pico kit on the Agilent 2100 Bioanalyzer System. Small RNA libraries were prepared from the isolated RNA using the Small RNA Library Prep Kit for Illumina, according to the manufacturer’s guidelines, as described in [29]. The quality of the libraries was checked using the High Sensitivity DNA Analysis Kit on the Bioanalyzer system before sequencing. Libraries were then denatured, diluted to a suitable concentration for optimal clustering, and sequenced on an Illumina NextSeq 500 platform. The size range specified for sequencing analysis was between 15-50 nucleotides (nt), covering the sizes of small RNAs and fragments from other RNAs with longer sizes. All procedures were performed at Norgen Biotek. Corp. (Thorold, Canada).

### Analysis of RNA sequencing data

#### Filtering, analysis, and gene counts

Raw sequence data were converted into fastq files and processed using the Genboree Workbench’s exceRpt small RNA-seq analysis pipeline (version 4.6.2, Extracellular RNA Communication Consortium – ERCC) [30]. Before sequence mapping to reference databases, adapters were trimmed, and reads shorter than 18 bp were removed. Low-quality sequences (with a Phred quality score below 30) and reads mapping to UniVec or human ribosomal RNA (rRNA) were also filtered out.

After data cleaning, the remaining reads were aligned to five reference databases (miRbase, gtRNAdb, piRNAbank, Gencode, and circBase), allowing for one maximum mismatch in each alignment. Raw count data was obtained for microRNAs (miRNA), transfer RNAs (tRNA), Piwi-interacting RNAs (piRNA), mitochondrial ribosomal RNAs (mt-rRNA), mitochondrial transfer RNAs (mt-tRNA), small nuclear RNAs (snRNA), small nucleolar RNAs (snoRNA), long intervening/intergenic noncoding RNAs (lincRNA), and circular RNAs (circRNA). For RNAs with a size longer than 50 nucleotides, read counts corresponded to fragments of the RNA biotype, but to simplify the same nomenclature was used for a fragment or a full RNA sequence. The quality of the libraries was continually monitored using FastQC software (version 0.11.9) during each step.

#### Differential expression analysis

Differential expression analysis was carried out using the DESeq2 package (version 1.34.0) for R, from Bioconductor [31]. Before count normalization, low expression RNAs were filtered out from analysis. We considered “low expression” when the sum of the RNA in all libraries did not reach a total of 1000 counts. mt-RNAs, snRNAs, and snoRNAs were excluded from normalization as these species exhibited read count patterns that were skewed towards the ataxic and pre-ataxic groups, with absolute expression levels differing substantially between the groups (see results for clarification). Such discrepancies would adversely affect the normalization of the overall dataset. Normalized counts were then calculated accounting for the depth of the libraries, using the DESeq2’s median of ratios normalization method. The negative binomial Wald test was used to model the data and to identify differentially expressed (DE) RNAs. The log fold-changes were shrunk using the adaptative shrinkage estimator, ‘ashr’, implemented elsewhere [32]. RNAs that matched the criteria of FDR-adjusted *p*-value < 0.05, and absolute log2 fold-change > 1 were considered DE. Heatmaps were created with the pheatmap package (version 1.0.12). Here, libraries and RNA biotypes were clustered using Ward’s minimum variance linkage method, and Euclidean distance was used as the dissimilarity measure. Principal component analyses (PCA) were performed using the plotPCA function from DESeq2, after variance stabilizing transformation of RNA-seq count data. Volcano plots were created with the EnhancedVolcano package (version 1.12.0) for R.

#### Gene ontology and pathway enrichment analysis

Functional over-representation analyses were performed using the subset of DE miRNAs between controls and ataxic mutation carriers. First, miRNAs were imported into the Mienturnet web platform [33], where potential target genes were predicted considering the miRTarBase database for miR-target interaction screening (minimum miR-target interactions = 5; FDR threshold ≤ 0.05). Subsequent analyses were performed using the clusterProfiler R package (version 4.2.2). The EnrichGO function was used to explore cellular components, molecular functions, and biological processes associated with the predicted target genes. In all cases, *p-values* were adjusted using the Benjamini-Hochberg correction.

### Statistical analysis

Continuous variables are reported as mean ± standard deviation. For inferential statistics, groups were compared using one-way analysis of variance (ANOVA) when variables displayed normal distribution, and Kruskal-Wallis test when a Gaussian distribution could not be assumed. Tukey’s and Dunn’s post-hoc tests were used to correct for multiple comparisons in the respective cases. Statistical significance was set at p < 0.05. To examine pairwise differences in miRNA, tRNA, and piRNA expression levels between control, pre-ataxic, and ataxic groups, we used the linear contrast in the DESeq2 package for RNA-seq differential analysis. For each pairwise comparison and for functional enrichment analysis, the false discovery rate (FDR) was controlled at 5 % using the Benjamini-Hochberg method. All statistical analyses were performed using GraphPad Prism version 9.3.1 (GraphPad Software, San Diego, USA) or R Statistical Software version 4.1.3 (R Core Team, 2022).

## Results

### Plasma-derived EVs isolated from ataxic and pre-ataxic SCA3 mutation carriers and controls are similar in size and concentration

We conducted an exploratory study to identify potential RNA biomarker candidates for SCA3 in plasma-derived EVs. A cohort of 32 individuals was divided into three groups: controls (n = 12), pre-ataxic mutation carriers (n = 10), and ataxic mutation carriers (n = 10; Figure 1). Although efforts were made to ensure a balanced distribution of age and gender, pre-ataxic mutation carriers were significantly younger than ataxic subjects (*p* < 0.01). The ataxic group showed an average SARA score of 18.0 ± 1.0, which is significantly higher than the pre-ataxic (*p* < 0.01) and the control group (*p* < 0.0001), as expected. The demographic and clinical information of the study participants is summarized in Table 1. EVs isolated from plasma samples showed no significant differences in size or concentration (particles/mL) between groups (Figure 2A-C). Yet, EVs from SCA3 mutation carriers tended to have lower RNA levels compared to control subjects (Figure 2D). The RNA yields obtained were consistent with other studies on plasma-derived EVs [34, 35] and met the minimum requirements for library preparation and sequencing.

**Figure 1.**
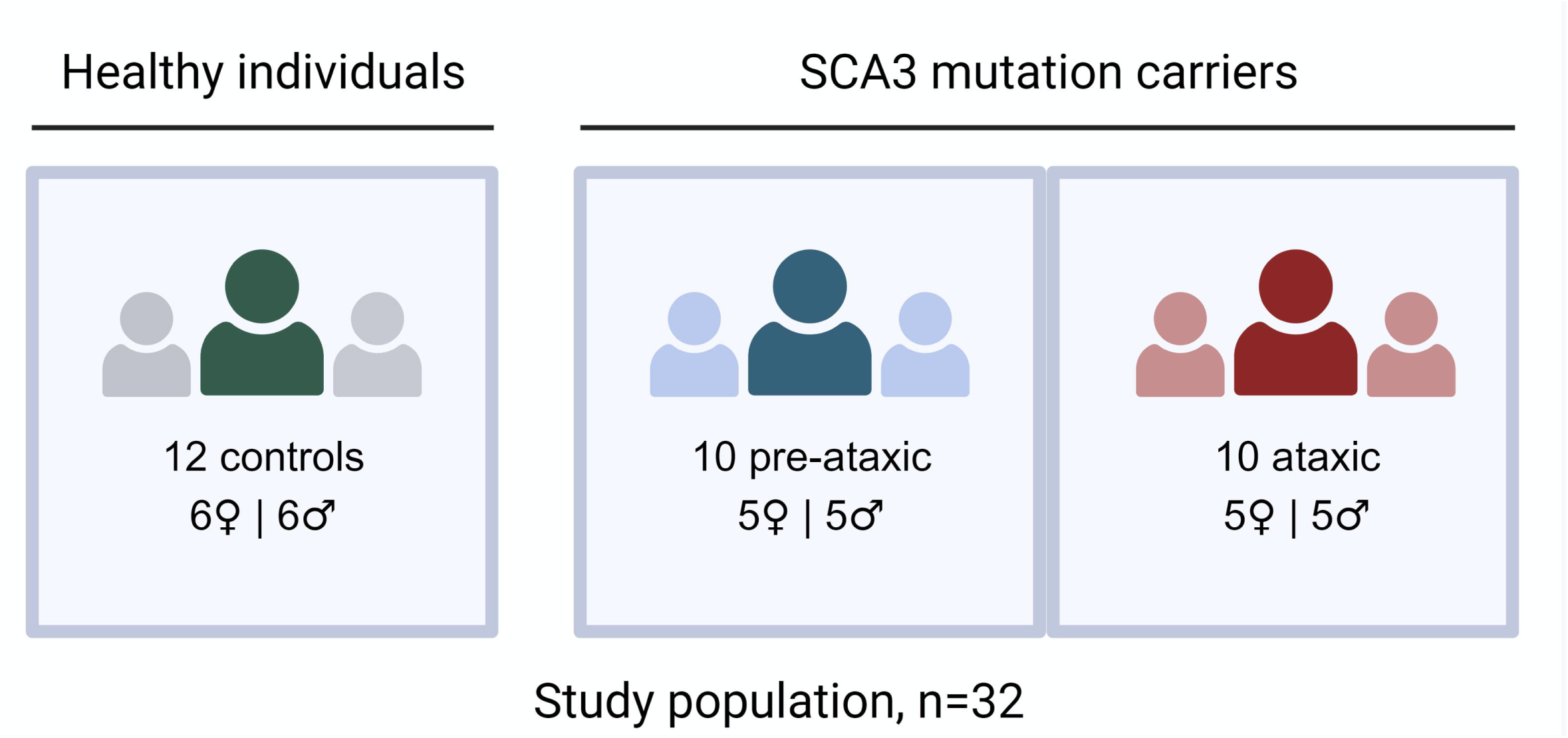
Study cohort composition. Study population consists of three distinct groups: healthy controls, SCA3 mutation carriers subdivided into two categories, pre-ataxic, and ataxic individuals. Each group was established to maintain a matching proportion of male and female individuals, ensuring gender balance within the study.

**Figure 2.**
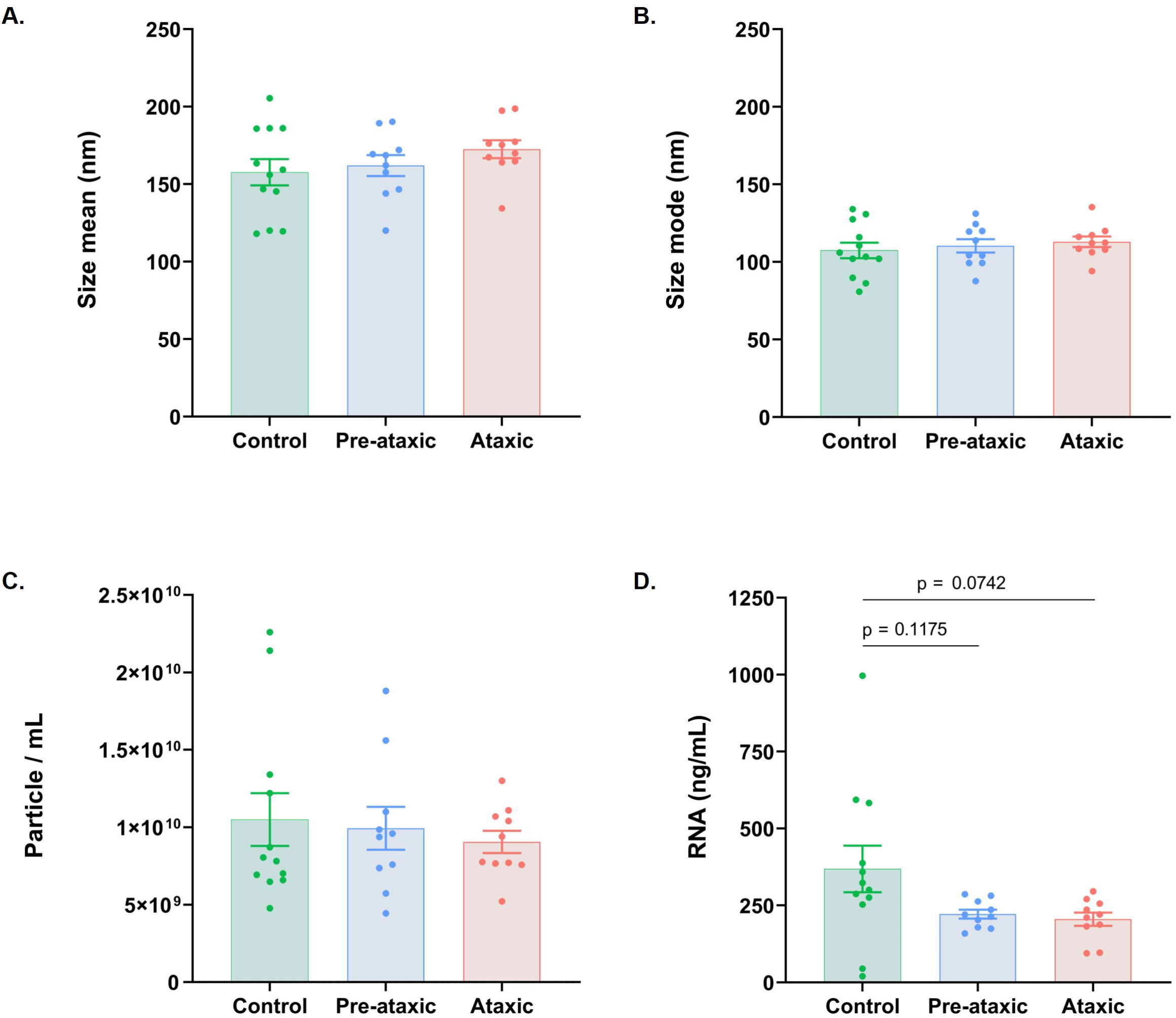
Size, particle concentration, and RNA content of plasma-derived EVs isolated from controls, pre-ataxic and ataxic SCA3 mutation carriers. The size and particle concentration of EVs from control (n = 12), pre-ataxic (n = 10) and ataxic mutation carriers (n = 10) were analyzed by nanoparticle tracking analysis (NTA). RNA was extracted, concentrated and quantified by Agilent RNA 6000 pico kit on the Agilent 2100 Bioanalyzer System. **(A)** EVs size (nm) presented as mean ± SEM. **(B)** EVs size (nm) presented as mode ± SEM. **(C)** EVs concentration (particle/mL) presented as mean ± SEM. **(D)** EVs RNA content (ng/mL) present as mean ± SEM. Statistical analysis was performed using one-way analysis of variance (ANOVA).

**Table 1.** Demographic and clinical characterization of the study population.

|  | Control<br>(C) | Pre-ataxic<br>(PA) | Ataxic<br>(A) | p-value<br>(C vs PA \| C vs A \| PA vs A) |
| --- | --- | --- | --- | --- |
| Sample size | 12 | 10 | 10 |  |
| Sex (m/f) | 6/6 | 5/5 | 5/5 | ns \| ns \| ns |
| Age | 41.5 [13.9] | 34.0 [6.65] | 51.3 [13.3] | ns \| ns \| ** |
| CAG | — | 67.8 [3.7] | 70.2 [4.3] | — \| — \| ns |
| SARA | 0.2 [0.6] | 0.8 [0.6] | 18.1 [1.2] | ns \| **** \| ** |

CAG: number of CAG triplet repeats; SARA: Scale for the Assessment and Rating of Ataxia. Data is presented as mean [ ± SD]. Statistics between groups were calculated by one-way ANOVA followed by a post-hoc Tukey's multiple comparison test (for age), unpaired Student's *t* test (for CAG), and Kruskal-Wallis followed by a post-hoc Dunn's multiple comparison test (for SARA). ns: non-significant ( $p>0.05$ ); \* $p<0.05$ ; \*\* $p<0.01$ ; \*\*\*\* $p<0.0001$ .

### Plasma-derived EVs from ataxic SCA3 mutation carriers are enriched in mitochondrial, nuclear, and nucleolar RNA biotypes compared to pre-ataxic and control subjects

In this study, the RNA sequencing focused on RNAs ranging from 15 to 50 nucleotides in length, covering most of the small RNAs and fragments from other RNAs biotypes with longer size. The obtained reads were trimmed, pre-processed for quality, and aligned and mapped to the human genome and reference RNA databases. The average input reads ranged from 7.7 to 44.7 million, with no significant differences between control (15.7 ± 6.0 million input reads), pre-ataxic (21.8 ± 11.6), and ataxic groups (14.0 ± 10.9). An average of 61.3 ± 4.7 % of the global input reads were clipped and subjected to a further quality control filter to exclude low-quality sequences, homopolymers, contaminants mapping to UniVec database, and ribosomal RNA (rRNA). Notably, a higher percentage of reads were filtered out in the ataxic group (11.9 ± 1.7 %) compared to control subjects (8.5 ± 0.9 %, *p* = 0.004) and pre-ataxic mutation carriers (8.8 ± 2.5 %, *p* = 0.001), likely due to the presence of higher rRNA levels in EVs of ataxic individuals (Supplementary Table 1). The remaining reads were mapped to the human genome and small RNA biotypes databases to study the small RNA content of plasma-derived EVs. The average number of reads mapping to the reference databases ranged from 1.6 to 2.6 million.

We then explored the abundance of RNA (or RNA fragments) mapping to the different biotypes between groups (Figure 3 and Table 2). Although the total number of RNA biotype counts was similar between the three groups, one participant in the control group (subject #10) was excluded from the analysis given the unique RNA biotype signature shown relative to all participants in study (controls and SCA3 mutation carriers; Supplementary Figure 1). MiRNAs were the most abundant biotype in all groups (81.4 ± 5.1 % of total mapped RNA), followed by tRNAs (5.7 ± 4.2 %) and piRNAs (4.3 ± 3.7 %). Protein-coding RNA fragments constituted up to 2.3 ± 1.0 % of the mapped small RNA in EVs. The least abundant biotypes were mt-rRNA, snRNA, mt-tRNA, snoRNA, and lincRNA, each accounting for less than 1 % of the total mapped RNA (Supplementary Figure 2). A miscellaneous group of other RNAs accounted for 3.2 ± 3.1 % of total RNAs in EVs. Compared to controls and pre-ataxic mutation carriers, EVs from ataxic subjects showed higher relative levels of mt-rRNAs (*p* = 0.0007), mt-tRNAs (*p* = 0.0003), snRNAs (*p* = 0.0003), and snoRNAs (*p* = 0.0003). Ataxic subjects also demonstrated a trend towards increased miRNA/tRNA ratios relative to controls (*p* = 0.0516). No other significant differences were found in the ratios of the three most prevalent RNA biotypes (miRNAs, piRNAs and tRNAs) between groups (Supplementary Table 2).

**Figure 3.**
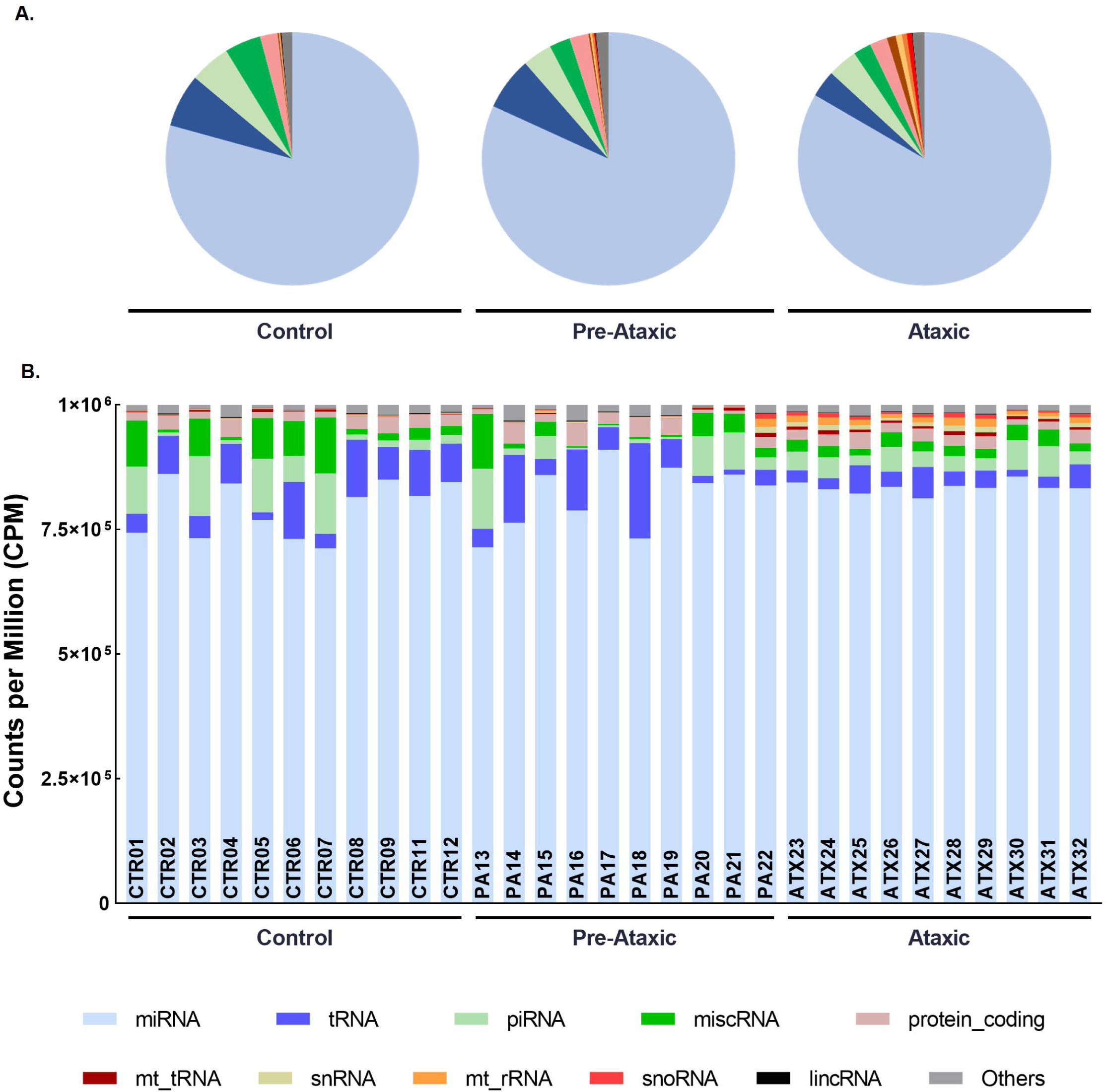
RNA biotype distribution in EVs from control, pre-ataxic and ataxic SCA3 mutation carriers. **(A)** A pie chart is presented for each group: healthy controls, pre-ataxic SCA3, and ataxic SCA3 mutation carriers. Each pie chart illustrates the distribution of RNA biotypes within the respective group. Each slice of the pie represents a specific RNA biotype, and the size of each slice corresponds to the proportion of that biotype within the group. **(B)** Bar plot representing the counts per million (CPM) of various RNA biotypes for each individual in the study cohort. Each bar corresponds to an individual, and the height of the bars reflects the CPM values of different RNA biotypes.

**Table 2.** Percentage of biotype counts in EVs of study groups.

|  |  | Control<br>(C) | Pre-ataxic<br>(PA) | Ataxic<br>(A) |  |
| --- | --- | --- | --- | --- | --- |
| Total million reads mapped to references |  | 1.59 [1.06] | 2.58 [1.48] | 1.97 [0.76] | p-value<br>(C vs PA \| C vs A \|PA vs A) |
| Biotype percentage (%) | miRNA | 79.3 [5.6] | 81.8 [6.5] | 83.4 [1.2] | ns \| ns \| ns |
|  | tRNA | 6.8 [3.3] | 6.8 [6.1] | 3.5 [1.6] | ns \| ns \| ns |
|  | piRNA | 5.2 [4.9] | 3.8 [4.1] | 3.8 [1.5] | ns \| ns \| ns |
|  | miscRNA | 4.6 [4.0] | 2.7 [3.3] | 2.3 [0.7] | ns \| ns \| ns |
|  | Protein coding RNA | 2.2 [0.8] | 2.4 [1.5] | 2.2 [0.6] | ns \| ns \| ns |
|  | mt_tRNA | 0.2 [0.2] | 0.2 [0.3] | 0.6 [0.1] | ns \| ** \| ** |
|  | snRNA | 0.1 [0.0] | 0.2 [0.3] | 0.8 [0.3] | ns \| *** \| ** |
|  | mt_rRNA | 0.1 [0.0] | 0.3 [0.5] | 1.2 [0.4] | ns \| *** \| ** |
|  | snoRNA | 0.0 [0.0] | 0.2 [0.3] | 0.6 [0.2] | ns \| *** \| ** |
|  | lincRNA | 0.1 [0.1] | 0.1 [0.1] | 0.1 [0.0] | ns \| ns \| ns |
|  | others | 1.4 [0.6] | 1.6 [1.1] | 1.5 [0.4] | ns \| ns \| ns |

Data is presented as mean [ ± SD]. Statistics between groups were calculated by one-way ANOVA followed by a post-hoc Tukey's multiple comparisons test. ns: non-significant ( $p>0.05$ ); \*\* $p<0.01$ ; \*\*\* $p<0.001$ ; \*\*\*\* $p<0.0001$ .

### Ataxic mutation carriers can be discriminated from control and pre-ataxic mutation carriers based on differentially expressed miRNAs

In previous studies, we and others have shown miRNA dysregulation in SCA3 [36–38]. Building on this, and considering the high abundance of miRNAs in EVs, we further investigated the miRNA content in plasma-derived EVs to identify potential biomarker candidates. A total of 1006 distinct miRNAs were identified. Among these, only 262 miRNAs (26.0% of the total) had an absolute cumulative read count above the established threshold of 1000 reads. The remaining 743 miRNAs, representing only 0.2% of the overall miRNA count, were excluded from subsequent analysis given their low expression levels (see Supplementary Tables 3.1 and 3.2 for details). All 262 miRNAs that passed the expression threshold were present in EVs of at least one participant from each group, indicating that none of the analyzed miRNAs was specific to any of the study groups. Among the 10 miRNAs with the highest abundance in each group, seven were common across all three groups (Table 3).

**Table 3.**
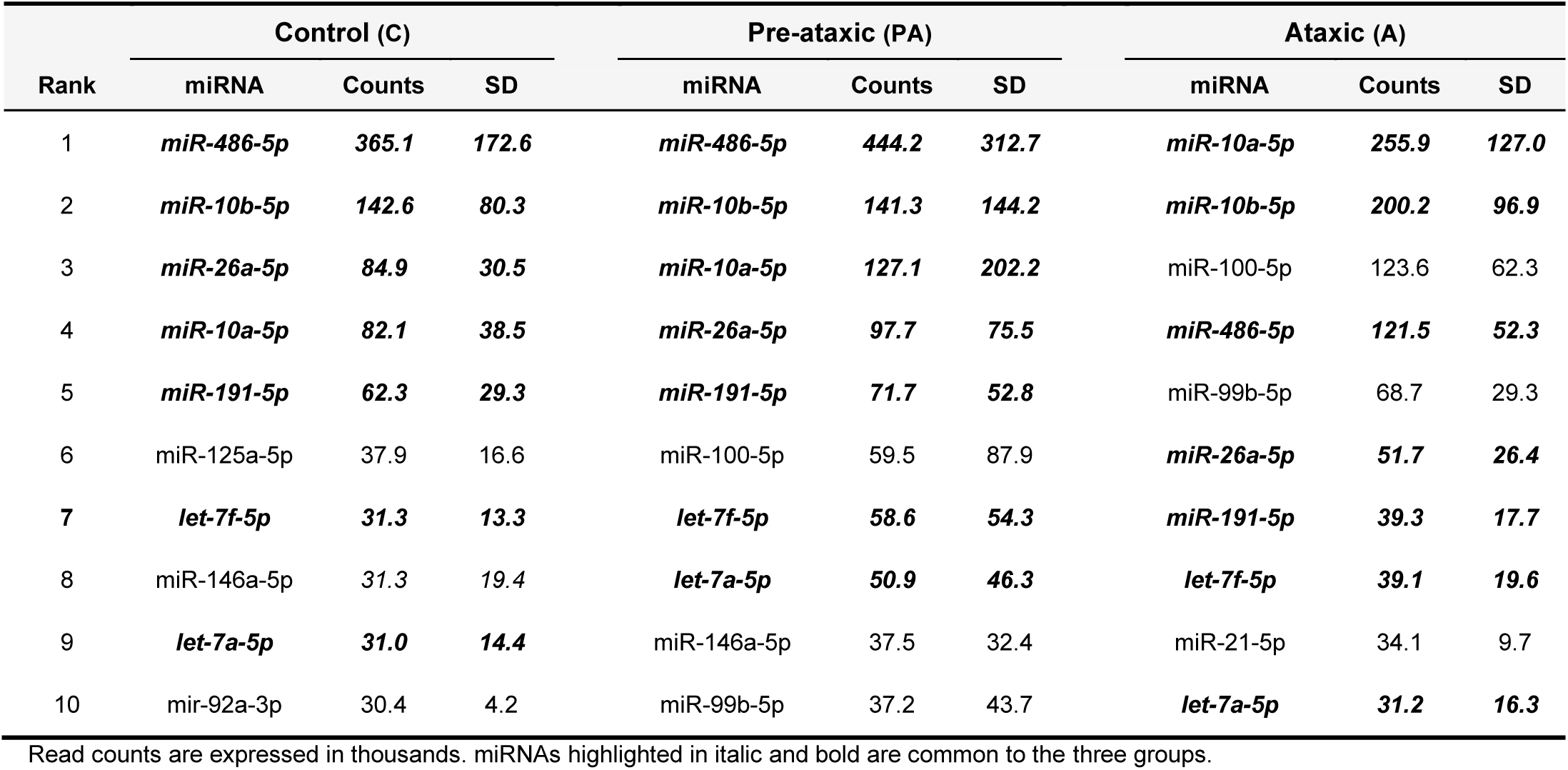
Average normalized read counts for the top 10 most abundant miRNAs in EVs of each study group.

Interestingly, when performing a principal component analysis (PCA) of the 262 miRNAs, ataxic mutation carriers were segregated from controls, and from pre-ataxic individuals, apart from one pre-ataxic subject that clustered among the ataxic group (Supplementary Figure 3). To determine which species were contributing the most to this segregation, we conducted a differential expression analysis (full results available in Supplementary Table 3.3). We found 53 miRNAs differentially expressed in ataxic mutation carriers compared to controls (criteria: absolute log_2_FC > 1; FDR < 0.05; Figure 4A and Table 4). Furthermore, 6 miRNAs were differentially expressed between pre-ataxic mutation carriers and controls, all overlapping with the former comparison (Figure 4B, 4D and Table 4). 38 miRNAs were differentially expressed between ataxic and pre-ataxic subjects, of which 25 overlapped with the first pairwise comparison (ataxic versus controls), while none overlapped with the second (pre-ataxic versus controls) (Figure 4C, 4D and Table 4).

**Figure 4.**
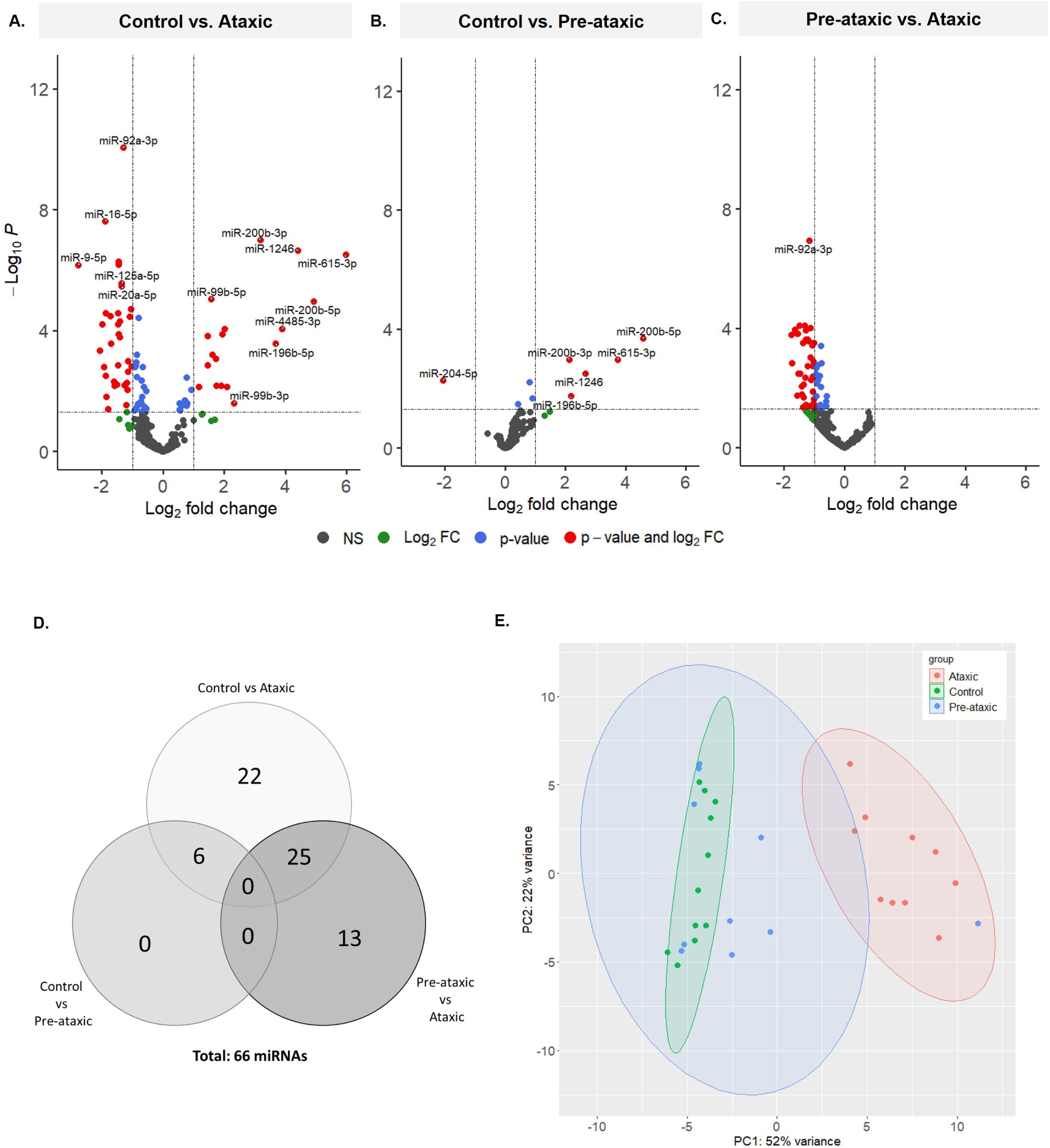
Differentially expressed miRNAs in EVs discriminate ataxic SCA3 mutation carriers from control and pre-ataxic subjects. **(A-C)** Volcano plots illustrating differentially expressed miRNAs between control and ataxic SCA3 mutation carriers (A), control and pre-ataxic SCA3 mutation carriers (B), and pre-ataxic and ataxic SCA3 mutation carriers (C). **(D)** Venn diagram of differentially expressed miRNAs identified in the three groups comparisons. **(E)** Principal component analysis of the differentially expressed miRNAs identified in groups comparisons (66 miRNAs corresponding to all red dots in volcano plots A and B).

**Table 4.** Top 30 differentially expressed miRNAs in EVs between the study groups, considering a FDR < 0.05 and an absolute Log2FC > 1.

| Control vs. Ataxic |  |  |  | Control vs. Pre-ataxic |  |  |  | Pre-ataxic vs. Ataxic |  |  |  |
| --- | --- | --- | --- | --- | --- | --- | --- | --- | --- | --- | --- |
| miRNA | baseMean | log2FC | FDR | miRNA | baseMean | log2FC | FDR | miRNA | baseMean | log2FC | FDR |
| miR-615-3p | 119 | 5.985 | <0.001 | miR-200b-5p | 50 | 4.585 | <0.001 | miR-144-3p | 5299 | -1.758 | <0.001 |
| miR-200b-5p | 50 | 4.932 | <0.001 | miR-615-3p | 119 | 3.732 | 0.001 | miR-9-5p | 144 | -1.733 | 0.001 |
| miR-1246 | 648 | 4.408 | <0.001 | miR-1246 | 648 | 2.657 | 0.003 | miR-486-5p | 312032 | -1.638 | <0.001 |
| miR-4485-3p | 243 | 3.891 | <0.001 | miR-196b-5p | 32 | 2.191 | 0.018 | miR-4732-3p | 60 | -1.575 | 0.018 |
| miR-196b-5p | 32 | 3.680 | <0.001 | miR-200b-3p | 2804 | 2.136 | 0.001 | miR-16-2-3p | 735 | -1.566 | <0.001 |
| miR-200b-3p | 2804 | 3.177 | <0.001 | miR-204-5p | 107 | -2.065 | 0.005 | miR-2110 | 242 | -1.554 | <0.001 |
| miR-9-5p | 144 | -2.778 | <0.001 |  |  |  |  | miR-144-5p | 1028 | -1.523 | 0.003 |
| miR-99b-3p | 59 | 2.324 | 0.025 |  |  |  |  | miR-16-5p | 2914 | -1.487 | <0.001 |
| miR-30c-2-3p | 89 | 2.090 | 0.007 |  |  |  |  | miR-451a | 20274 | -1.465 | 0.003 |
| miR-124-3p | 356 | -2.075 | <0.001 |  |  |  |  | miR-124-3p | 356 | -1.410 | 0.017 |
| miR-100-5p | 68548 | 2.015 | <0.001 |  |  |  |  | miR-127-3p | 2245 | -1.406 | 0.009 |
| miR-144-5p | 1028 | -1.989 | <0.001 |  |  |  |  | miR-363-3p | 1766 | -1.386 | <0.001 |
| miR-30a-3p | 1442 | 1.941 | <0.001 |  |  |  |  | miR-3158-3p | 42 | -1.370 | 0.043 |
| miR-29b-3p | 122 | -1.938 | 0.002 |  |  |  |  | miR-142-3p | 151 | -1.354 | 0.020 |
| miR-21-3p | 439 | 1.899 | 0.007 |  |  |  |  | miR-1-3p | 798 | -1.352 | 0.008 |
| miR-16-5p | 2914 | -1.889 | <0.001 |  |  |  |  | miR-7-5p | 224 | -1.314 | <0.001 |
| miR-144-3p | 5299 | -1.874 | <0.001 |  |  |  |  | miR-150-5p | 2833 | -1.294 | 0.004 |
| miR-204-5p | 107 | -1.868 | 0.003 |  |  |  |  | miR-29b-3p | 122 | -1.288 | 0.037 |
| miR-29c-3p | 65 | -1.846 | 0.015 |  |  |  |  | miR-143-3p | 6175 | -1.284 | <0.001 |
| miR-3158-3p | 42 | -1.799 | 0.039 |  |  |  |  | miR-101-3p | 5467 | -1.274 | <0.001 |
| miR-200a-3p | 219 | 1.755 | 0.007 |  |  |  |  | miR-20a-5p | 585 | -1.225 | <0.001 |
| miR-183-5p | 7219 | 1.734 | 0.001 |  |  |  |  | miR-142-5p | 2167 | -1.222 | 0.002 |
| miR-150-5p | 2833 | -1.724 | <0.001 |  |  |  |  | miR-484 | 3121 | -1.180 | <0.001 |
| miR-451a | 20274 | -1.700 | <0.001 |  |  |  |  | miR-150-3p | 130 | -1.173 | 0.040 |
| miR-30a-5p | 18689 | 1.618 | 0.001 |  |  |  |  | miR-92a-3p | 23567 | -1.159 | <0.001 |
| miR-142-3p | 151 | -1.603 | 0.005 |  |  |  |  | miR-423-5p | 19369 | -1.130 | 0.001 |
| miR-215-5p | 47 | -1.591 | 0.007 |  |  |  |  | miR-185-5p | 3500 | -1.122 | <0.001 |
| miR-99b-5p | 41802 | 1.567 | <0.001 |  |  |  |  | miR-126-5p | 5064 | -1.111 | 0.005 |
| miR-122-5p | 5805 | -1.511 | 0.006 |  |  |  |  | miR-186-5p | 4397 | -1.064 | <0.001 |
| miR-126-5p | 5064 | -1.476 | <0.001 |  |  |  |  | miR-30e-5p | 4944 | -1.056 | 0.004 |

Data is sorted by absolute log2FC from highest to lowest. baseMean: average normalized counts; log2FC: log2 fold-change between the groups; FDR: Benjamini-Hochberg adjusted p-value.

Altogether, the three pairwise comparisons (control versus ataxic, control versus pre-ataxic, and pre-ataxic versus ataxic) resulted in 66 unique differentially expressed miRNAs (Figure 4D). This panel of differentially expressed miRNAs was further used to investigate whether this smaller subset could also discriminate ataxic SCA3 mutation carriers from pre-ataxic and controls. The results, presented in Figure 4E, demonstrate that this panel of miRNAs successfully separated ataxic mutation carriers from pre-ataxic carriers and control subjects, except for one pre-ataxic subject that was not distinguished from the ataxic group cluster. This finding is supported by a heatmap analysis of the same 66 miRNAs, which clearly shows a hierarchical grouping of the ataxic subjects (Supplementary Figure 4). Control subjects and pre-ataxic mutation carriers were not distinguished whether using all 262 miRNAs (Supplementary Figure 3) or the 66 differentially expressed miRNAs (Figure 4E).

### The target genes of differentially expressed miRNAs are involved in stress response, tissue development, regulation of protein stability and miRNA processes

To uncover the regulatory mechanisms of the miRNAs found to be differentially expressed in EVs of SCA3 ataxic mutation carriers and identify potentially altered processes in the disease, we performed a functional over-representation analysis (ORA) using the target genes of the differentially expressed miRNAs between ataxic mutation carriers and control subjects. Such target genes were predicted using the Mienturnet web platform (the criteria for miRNA-target interaction screening consisted of a minimum of 5 interactions and an FDR < 0.05). A total of 454 potential target genes were identified using miRTarBase database (Supplementary Table 4). Based on these, we further performed Gene Ontology (GO) functional over-representation analysis for biological processes, molecular functions, and cellular components. Our results showed an enrichment of biological processes related to cellular responses to oxidative/chemical stress and low oxygen levels, forebrain and gland development, tissue migration, and regulation of protein stability and miRNA processes (metabolic and transcription) (Figure 5A). Moreover, we observed an enrichment in molecular functions related to DNA-binding (RNA polymerase II-specific DNA-binding transcription factor binding, DNA-binding transcription activator activity, transcription coregulator binding and chromatin DNA binding), as well as protein binding (cadherin, ubiquitin protein ligase, phosphatase, β-catenin) (Figure 5B). We also found that cell-substrate junctions, cell leading edge, nuclear membrane and RNA polymerase II transcription regulator complex are amongst the cellular components mostly associated with the 454 target genes (Figure 5C).

**Figure 5.**
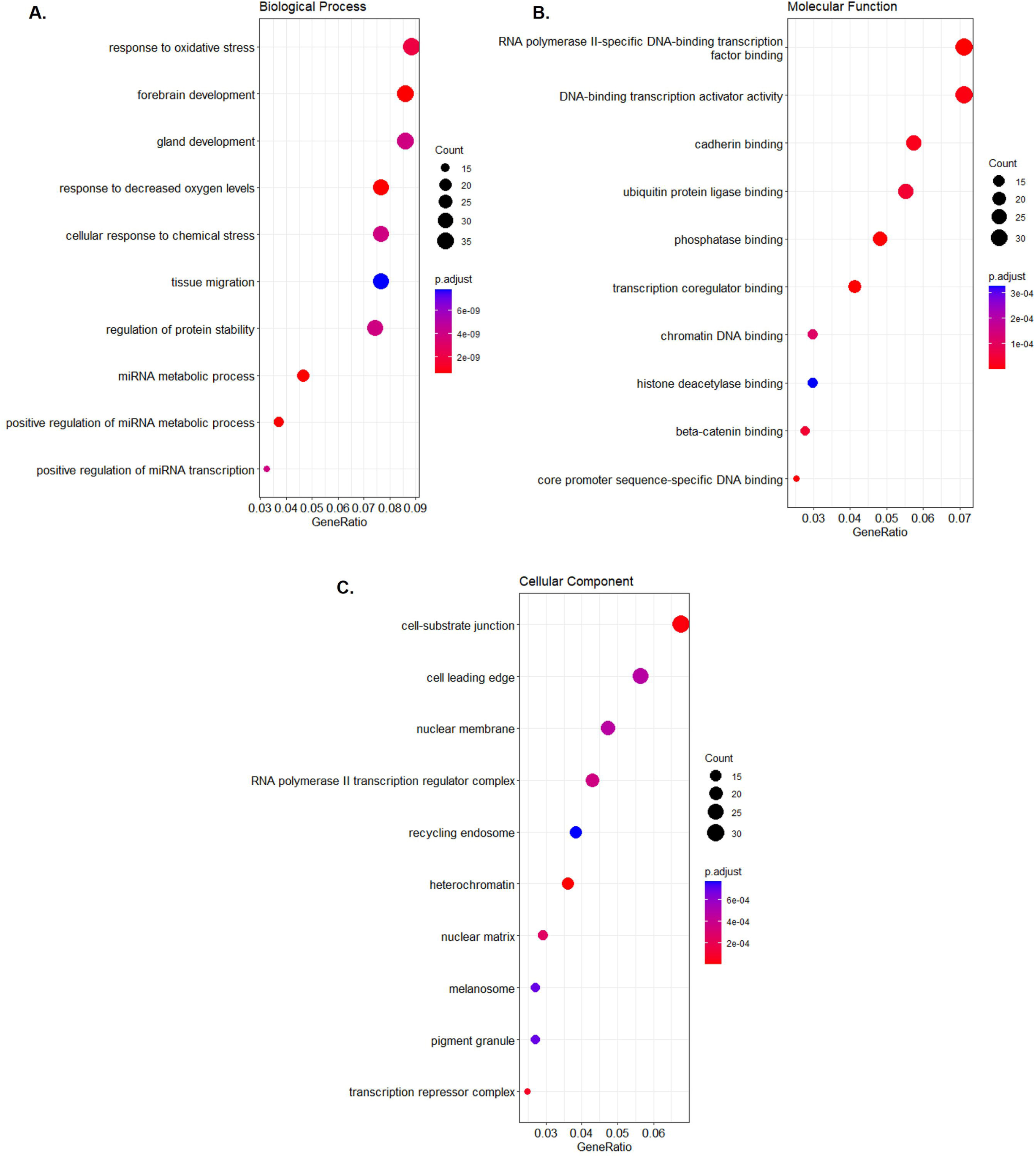
Gene Ontology (GO) enrichment analysis of target genes of DE miRNAs. The top ten enriched GO biological processes **(A)**, molecular functions (**B)**, and cellular components **(C)** for the identified targets of the DE miRNAs between controls and ataxic mutation carriers. The color and size of the bubbles in the figure correspond to FDR values and number of interacting target genes, respectively.

### Twelve piRNAs can effectively distinguish ataxic SCA3 mutation carriers from pre-ataxic and control subjects

We also found tRNAs and piRNAs to be present with considerable abundance in plasma-EVs of the study subjects. Hence, we explored if these RNAs species were also able to distinguish mutation carriers from control subjects.

Regarding tRNAs, all 23 domain-specific tRNAs were detected in all sample libraries, including the tRNA^iMet^ (responsible for carrying the amino acid methionine (Met) to the ribosome during protein synthesis; the “i” stands for “initiator”, as tRNA^iMet^ recognizes the start codon AUG, the codon that initiates protein synthesis [39]) and the suppressor-tRNA. Among these, only tRNA^Asn^ failed to reach the minimum threshold of 1000 read count across libraries and was excluded from further analysis. Globally, the most abundant domain-specific tRNAs were tRNA^Gly^, tRNA^Lys^, tRNA^Glu^, tRNA^Ala^, and tRNA^Cys^ (Supplementary Tables 5.1 and 5.2). Differential expression analysis revealed significant differences (absolute log_2_FC>1) between ataxic and control subjects in 6 out of the 22 tRNAs. Among these, tRNA^Gly^ and tRNA^Ile^ were also found to be significantly downregulated in EVs from ataxic mutation carriers compared to pre-ataxic mutation carriers, whereas no tRNA was found differentially expressed between control and pre-ataxic groups (Figure 6; Table 5; Supplementary Table 5.3). PCA analysis of both the entire set of 22 tRNAs (Supplementary Figure 5) and the subset of 6 differentially expressed tRNAs (Figure 6D) failed to efficiently segregate the three study groups.

**Figure 6.**
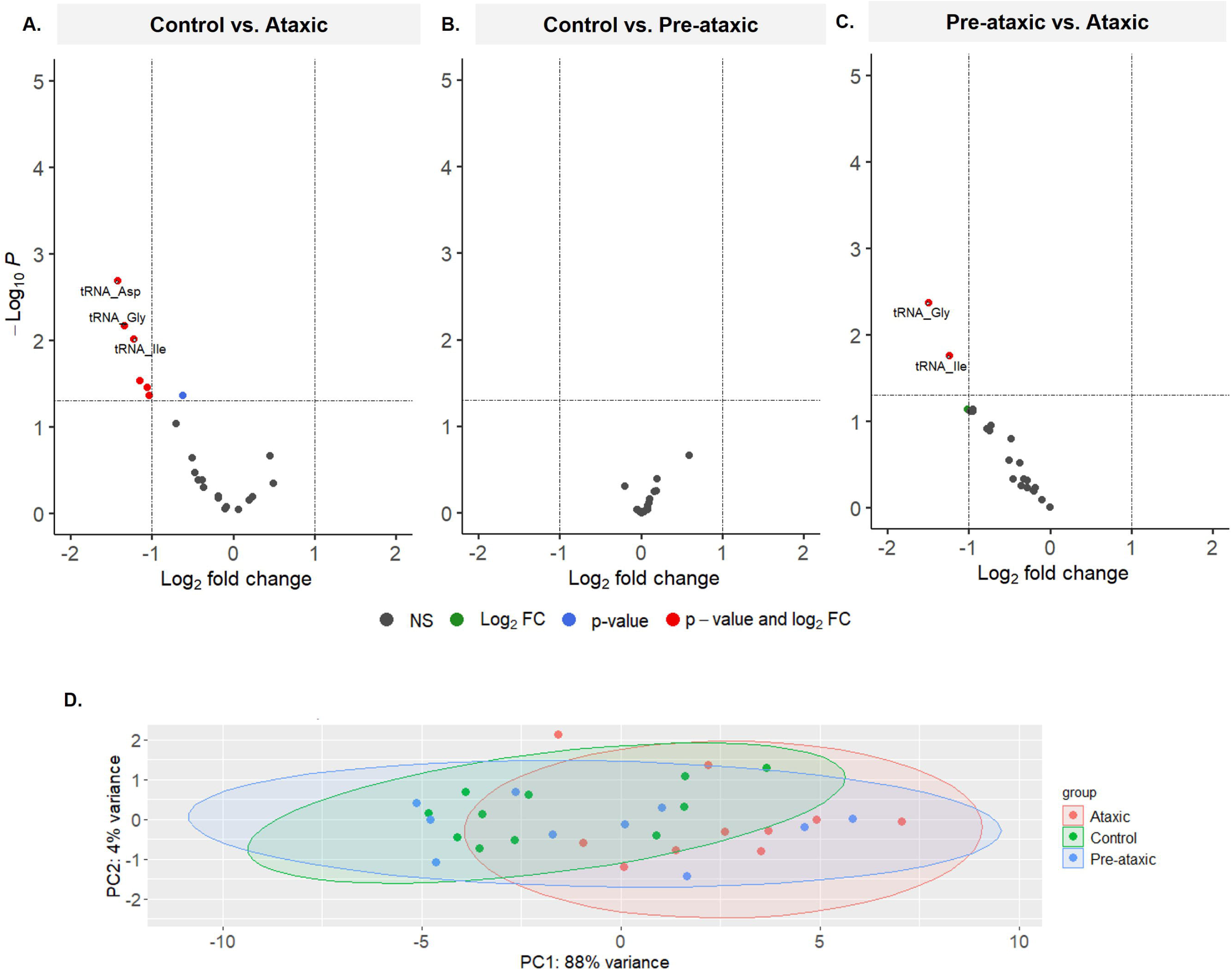
Differentially expressed tRNAs in EVs of ataxic SCA3 mutation carriers from control and pre-ataxic subjects. **(A-C)** Volcano plots illustrating differentially expressed tRNAs between control and ataxic SCA3 mutation carriers (A), control and pre-ataxic SCA3 mutation carriers (B), and pre-ataxic and ataxic SCA3 mutation carriers (C). **(D)** Principal component analysis of the 6 differentially expressed tRNAs identified in groups comparisons.

**Table 5.** Differentially expressed tRNAs, considering a FDR < 0.05 and an absolute Log2FC > 1.

| Control vs. Ataxic |  |  |  | Pre-ataxic vs. Ataxic |  |  |  |
| --- | --- | --- | --- | --- | --- | --- | --- |
| RNA | baseMean | log2FC | FDR | RNA | baseMean | log2FC | FDR |
| tRNA_Asp | 284 | -1.422 | 0.002 | tRNA_Gly | 66269 | -1.495 | 0.004 |
| tRNA_Gly | 66269 | -1.341 | 0.007 | tRNA_Ile | 297 | -1.244 | 0.017 |
| tRNA_Ile | 297 | -1.223 | 0.010 |  |  |  |  |
| tRNA_Sup | 143 | -1.150 | 0.029 |  |  |  |  |
| tRNA_Phe | 222 | -1.062 | 0.035 |  |  |  |  |
| tRNA_Cys | 5245 | -1.035 | 0.043 |  |  |  |  |

Data is sorted by absolute log2FC from highest to lowest. baseMean: average normalized counts; log2FC: log2 fold-change between the groups; FDR: Benjamini-Hochberg adjusted p-value.

Concerning piRNAs, we found 157 different piRNA species present in EV. Yet, the majority of these piRNA species were present in low levels. Out of the 157 piRNA species, only 20 showed an absolute read count above the threshold of 1000 and were therefore considered for differential expression analysis (Supplementary Tables 6.1 and 6.2). The most abundant piRNAs in all three groups were piR_016658 and piR_016659. A differential expression analysis revealed 12 differentially expressed piRNAs in plasma-derived EVs from ataxic mutation carriers compared to controls. Six of these were also differentially expressed in the control vs pre-ataxic group pairwise comparison (Table 6; Figure 7A-B; Supplementary Table 6.3). Finally, piR_019825 and piR_016659 showed significantly lower levels in EVs from ataxic compared to those obtained from pre-ataxic mutation carriers (Figure 7C; Table 6; Supplementary Table 6.3). Again, we performed PCA analyses to investigate if it was possible to cluster and distinguish the study subjects based on the expression levels of the entire set of piRNAs, as well as the subset of differentially expressed piRNAs. The obtained results demonstrated that both the entire set of piRNAs (Supplementary Figure 6) and the differentially expressed subset (Figure 7D) could distinguish the ataxic group from the remaining individuals (except for one pre-ataxic mutation carrier). Remarkably, principal component 1 (PC1), the one most contributing for the segregation of the clusters, explained over 78% of the variation in the data (Figure 7D).

**Figure 7.**
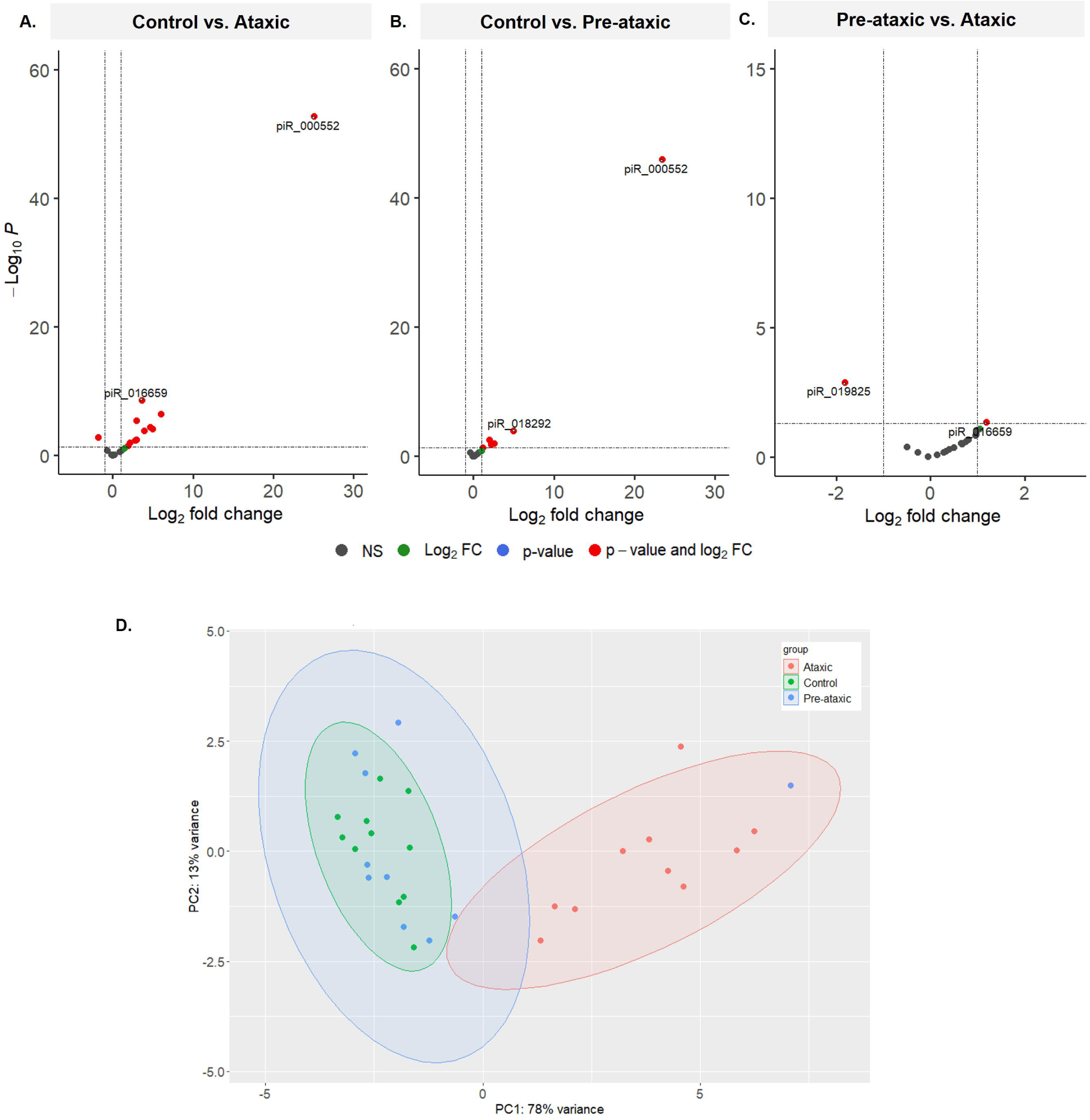
Differentially expressed piRNAs in EVs discriminate ataxic SCA3 mutation carriers from control and pre-ataxic subjects. **(A-C)** Volcano plots illustrating differentially expressed piRNAs between control and ataxic SCA3 mutation carriers (A), control and pre-ataxic SCA3 mutation carriers (B), and pre-ataxic and ataxic SCA3 mutation carriers (C). **(D)** Principal component analysis of the 12 differentially expressed piRNAs identified in groups comparisons.

**Table 6.** Differentially expressed piRNAs, considering a FDR < 0.05 and an absolute Log2FC > 1.

| Control vs. Ataxic |  |  |  | Control vs. Pre-ataxic |  |  |  | Pre-ataxic vs. Ataxic |  |  |  |
| --- | --- | --- | --- | --- | --- | --- | --- | --- | --- | --- | --- |
| RNA | baseMean | log2FC | FDR | RNA | baseMean | log2FC | FDR | RNA | baseMean | log2FC | FDR |
| piR_000552 | 68 | 25.055 | <0.001 | piR_000552 | 68 | 23.434 | <0.001 | piR_019825 | 546 | -1.817 | 0.001 |
| piR_018292 | 30 | 5.970 | <0.001 | piR_018292 | 30 | 4.944 | <0.001 | piR_016659 | 5407 | 1.180 | 0.045 |
| piR_020619 | 30 | 4.992 | <0.001 | piR_020619 | 30 | 2.570 | 0.010 |  |  |  |  |
| piR_017184 | 72 | 4.638 | <0.001 | piR_017184 | 72 | 2.212 | 0.018 |  |  |  |  |
| piR_001170 | 29 | 3.930 | <0.001 | piR_016659 | 5407 | 1.991 | 0.003 |  |  |  |  |
| piR_016659 | 5407 | 3.601 | <0.001 | piR_014620 | 487 | 1.182 | 0.050 |  |  |  |  |
| piR_014620 | 487 | 2.958 | <0.001 |  |  |  |  |  |  |  |  |
| piR_020828 | 33 | 2.942 | 0.004 |  |  |  |  |  |  |  |  |
| piR_020008 | 56 | 2.797 | 0.005 |  |  |  |  |  |  |  |  |
| piR_019521 | 140 | 2.143 | 0.010 |  |  |  |  |  |  |  |  |
| piR_020450 | 115 | 1.908 | 0.030 |  |  |  |  |  |  |  |  |
| piR_019825 | 546 | -1.848 | 0.002 |  |  |  |  |  |  |  |  |

Data is sorted by absolute log2FC from highest to lowest. baseMean: average normalized counts; log2FC: log2 fold-change between the groups; FDR: Benjamini-Hochberg adjusted p-value.

## Discussion

In this study, we analyzed the small non-coding RNA content of plasma-derived EVs of both pre-ataxic and ataxic SCA3 mutation carriers, compared to control subjects. Our results show evident differences in the abundance of specific RNA biotypes and levels of specific small RNAs mostly between SCA3 ataxic patients and control subjects. We report for the first time an enrichment of mitochondrial, nuclear, and nucleolar RNA biotypes in EVs from SCA3 ataxic patients. We observed that the EVs content in some small specific RNA species, particularly miRNA and piRNAs, could distinguish ataxic SCA3 mutation carriers from controls and pre-ataxic individuals.

EVs are very attractive for biomarker research due to their cargo that may reveal unique signatures of pathological states [22]. Widespread employment of RNA sequencing has demonstrated that EVs can contain many RNA biotypes [40]. Among the small non-coding RNAs, miRNAs have become the most explored class, likely given their known role in gene regulation and their abundance in vesicles [40]. Altered miRNAs profiles have already been reported in SCA3 in patient-derived cells [36, 41, 42], in animal models [36], and in peripheral samples of patients [37, 38]. MiRNA dysregulation in SCA3 is corroborated in this study as we show that ataxic SCA3 mutation carriers can be discriminated from controls and pre-ataxic mutation carriers based on the miRNA content of plasma-derived EVs. This observation suggests that alterations in the miRNA profile are broad and occur with disease progression. Amongst the identified differentially expressed miRNAs in our study, only miR-9-5p has been previously linked to SCA3. miR-9 is predicted to target the 3’UTR of *ATXN3.* Carmona et al. found that the expression levels of miR-9 were decreased in SCA3 rodent models and post-mortem human brain samples [36]. In our study, we also observed lower levels of miR-9-5p in plasma-derived EVs of patients with SCA3, relative to control subjects. These observations are consistent with findings from other independent studies in various neurodegenerative diseases, such as Alzheimer’s disease [43, 44], amyotrophic lateral sclerosis [45], and Huntington’s disease [46].

Hou and colleagues have also recently explored the miRNA content of plasma-derived EVs from ataxic SCA3 mutation carriers, relative to control subjects [38]. The authors report several differentially expressed miRNAs, however, only 6 of these DE miRNAs overlap with those we identified in our study (miR-122-5p, miR-142-3p, miR-146b-5p, miR-194-5p, miR-196b-5p, miR-200a-3p, miR-204-5p and miR-486-3p) and only 4 were altered in the same direction (miR-122-5p, miR-194-5p, miR-196b-5p and miR-200a-3p). Technical differences between studies have often resulted in conflicting data due to differences in isolation of EV sub-populations and/or contaminants [47]. Indeed, while we used SEC for EVs isolation, Hou and colleagues employed a commercial kit based on the principle of precipitation. While SEC is considered a method of intermediate recovery and specificity, precipitation methods have low specificity and co-isolate several contaminants present in plasma [48, 49]. Discrepancies between studies are further exacerbated by other factors, such as differences in cohort size and study participants, sequencing platforms and techniques, and statistical methods for DE analysis.

We also investigated the predicted downstream targets of the identified differentially expressed miRNAs to explore and better understand altered processes in SCA3. The most enriched biological processes were associated to cellular responses to oxidative/chemical stress, tissue development and migration, and regulation of protein stability and miRNA processes. Most of these biological processes have been associated with the pathogenesis of SCA3 [50, 51]. This suggests that the identified dysregulated miRNAs are involved in and might contribute to mechanisms underlying the disease, and therefore may constitute potential therapeutic targets.

More recently, other classes of RNAs, namely tRNAs, piRNAs, snoRNAs, and mtRNAs, were also shown to be dysregulated in the context of neurodegenerative diseases [52–55]. tRNAs and piRNAs revealed to be the second and third most abundant RNA biotypes in EVs across all groups, respectively. While the biological significance of specific tRNAs identified as differentially expressed in ataxic mutation carriers might be questionable due to their low fold-change, the differences observed in the levels of certain piRNAs suggest that these may play an important role in the pathogenesis of the disease. piRNAs are known to regulate gene expression by suppressing transposable DNA elements that can integrate into new positions in the genome [56]. We found that the expression levels of a panel of 20 piRNAs (all that surpassed the threshold of 1000 reads across all libraries) is sufficient to discriminate ataxic SCA3 patients from pre-ataxic and control subjects. Similar results were achieved when we condensed the analysis to the 12 DE piRNAs. Dysregulation of piRNAs have already been observed in other neurodegenerative disorders [57, 58], but the exact mechanisms underlying this dysregulation and its consequences in these conditions remain unclear and require further investigation. To the best of our knowledge, no prior studies have explored the putative role of piRNAs in the context of SCA3.

Finally, an unexpected finding from our study was related to less abundant RNA biotypes. Specifically, we discovered that EVs from ataxic SCA3 patients have a significant enrichment of mtRNA (mt-rRNA and mt-tRNA), snRNAs, and snoRNAs compared to the other groups. While these biotypes were barely detected in EVs from controls and pre-ataxic mutation carriers (less than 1 % of the global small RNA content in both groups), the levels of these RNA biotypes were significantly higher in EVs from ataxic SCA3 mutation carriers. Recent evidence has shown that specific mtRNAs, snRNAs and snoRNAs are differentially expressed in EVs of patients with Alzheimer’s disease [53, 59]. However, the role of these RNA species in disease mechanisms remains largely unknown. While we might speculate that mitochondrial dysfunction in neurodegenerative diseases may lead to the release of mtRNA into the cytosol [60] and its incorporation in EVs, the mechanisms underlying the enrichment in snoRNAs and snRNAs in EVs, and their relation to disease pathogenesis, require further investigation.

The small sample size and the lack of validation of our findings in a larger cohort are the major limitations of this study. Validation of RNA sequencing results by the less expensive real-time quantitative PCR (qPCR) technique has been the most widely used for RNA biomarker studies [61]. However, qPCR is highly dependent on the normalization with a reference gene and does not allow the determination of RNA biotype abundance, which according to our results might itself be useful as a biomarker. Thus, we consider our findings would benefit from a validation approach in a large set of samples but using an RNA sequencing platform. The decreasing costs of RNA sequencing will allow a cost-effective strategy to validate these findings in a near future [61].

In summary, our study provides a comprehensive analysis of the small RNA biotypes present in plasma-derived EVs of ataxic and pre-ataxic SCA3 mutation carriers, and control subjects. Our findings reveal that EVs derived from ataxic individuals are enriched in mt-rRNA, mt-tRNA, snoRNA, and snRNA. Furthermore, we identified subsets of differentially expressed RNAs for the most abundant biotypes - miRNAs, tRNA and piRNA that might be promising biomarker candidates for SCA3. Remarkably, our study demonstrates that miRNA and piRNAs content in EVs can effectively distinguish ataxic subjects from the controls and pre-ataxic mutation carriers. Most of the differentially expressed RNAs reported in our study have never been associated with SCA3 or any other neurodegenerative disorder. These findings provide an opportunity for more focused studies to understand their role in disease pathogenesis and explore their potential as disease biomarkers, which ultimately may unveil novel therapeutic targets.

## Acknowledgements

The ESMI consortium acknowledges Ruth Hossinger for the project management of the ESMI project and for all contributions made towards the success of this project.

## Funding

This publication is an outcome of ESMI, an EU Joint Programme - Neurodegenerative Disease Research (JPND) project (see www.jpnd.eu). The project is supported through the following funding organisations under the aegis of JPND: Germany, Federal Ministry of Education and Research (BMBF; funding codes 01ED1602A/B); Netherlands, The Netherlands Organisation for Health Research and Development; Portugal, Fundação para a Ciência e Tecnologia (funding code JPCOFUND/0002/2015); United Kingdom, Medical Research Council (MR/N028767/1). This project has received funding from the European Union’s Horizon 2020 research and innovation programme under grant agreement No 643417. At the Portuguese sites, LPA also received funding from European Regional Development Fund (ERDF), through the Centro 2020 Regional Operational Program; through the COMPETE 2020 - Operational Programme for Competitiveness and Internationalisation, and Portuguese national funds via FCT – Fundação para a Ciência e a Tecnologia, under the projects: CENTRO-01-0145-FEDER-181240, UIDB/04539/2020, UIDP/04539/2020, LA/P/0058/2020, ViraVector (CENTRO-01-0145-FEDER-022095), BDforMJD (CENTRO-01-0145-FEDER-181240 & 2022.06118.PTDC), ARDAT under the IMI2 JU Grant agreement No 945473 supported by the European Union’s H2020 programme and EFPIA; by the American Portuguese Biomedical Research Fund (APBRF), National Ataxia Foundation and the Richard Chin and Lily Lock Machado-Joseph Disease Research Fund.

On the Azores, ESMI Network is currently supported by the Regional Government (Fundo Regional para a Ciência e a Tecnologia-FRCT), under the PRO-SCIENTIA program.

The work by MSyn was also supported by the JPND grant “GENFI-prox” (by DLR/BMBF).

PG is supported by the National Institute for Health Research University College London Hospitals Biomedical Research Centre UCLH. PG receives also support from the North Thames CRN.

BvdW receives funding from ZonMw, NWO, Gossweiler Foundation, Brugling Fonds, Radboudumc, Christina Foundation and Hersenstichting.

## Competing interests

TK, MS, BvdW are members of the European Reference Network for Rare Neurological Diseases (ERN-RD, project number 739510). PS and MMP were supported by FCT under the fellowship grants SFRH/BD/148451/2019 and 2022.11089.BD, respectively. TMM was supported by a fellowship from the BioSys PhD programme PD65-2012 (PD/BD/142854/2018) also from FCT. MR is supported by FCT CEECIND/03018/2018. PG and HGM work at University College London Hospitals/University College London, which receives a proportion of funding from the Department of Health’s National Institute for Health Research Biomedical Research Centre’s funding scheme. PG received funding from the Medical Research Council (MR/N028767/1) and CureSCA3 in support of HGM work. JF received funding as a fellow of the Hertie Network of Excellence in Clinical Neuroscience.

MS has received consultancy honoraria from Janssen, Ionis, Orphazyme, Servier, Reata, GenOrph, and AviadoBio, all unrelated to the present manuscript. LPA research group has received private funding from PTC Therapeutics, Uniqure, Wave life Sciences, Servier, Blade Therapeutics and Hoffmann-La Roche AG outside the submitted work. Radboud University Medical Center (through BvdW) has received consultancy fees from Servier and Biohaven, and BvdW serves on a scientific advisory board of Vico Therapeutics.

PG has received grants and honoraria for advisory board from Vico Therapeutics, honoraria for advisory board from Triplet Therapeutics, grants and personal fees from Reata Pharmaceutical, grants from Wave.

## Author’s contribution

MMS, PS: study concept and design, sample collection and preparation, data acquisition, experimental execution, data analysis and discussion, manuscript writing and revision

MMP, LG: sample collection, data acquisition, data analysis and discussion, manuscript review

RN, SPD, TMM, MGC: data analysis and discussion

CJ, JR, IC, JHS, JI: data acquisition and discussion

JHS, JI, MR, HGM: sample collection, data acquisition and manuscript review

JF, MS: data acquisition and manuscript review

MS, ML, PG, BvdW, TK: study concept and design, and manuscript review

LPA: Study concept and design, data discussion, manuscript review

## Appendix 1. Additional members of the European Spinocerebellar Ataxia type 3/Machado-Joseph Initiative (ESMI) Study Group

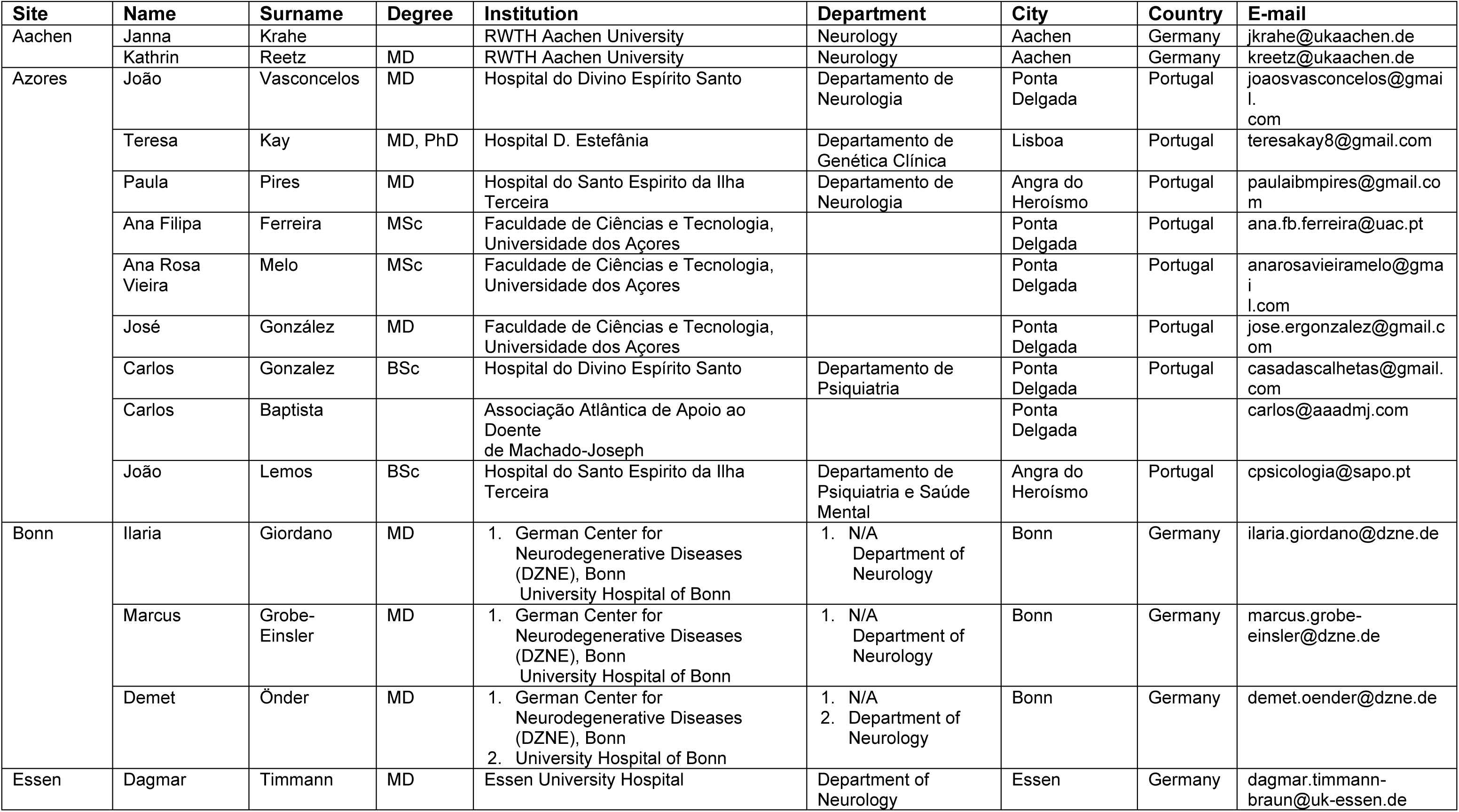
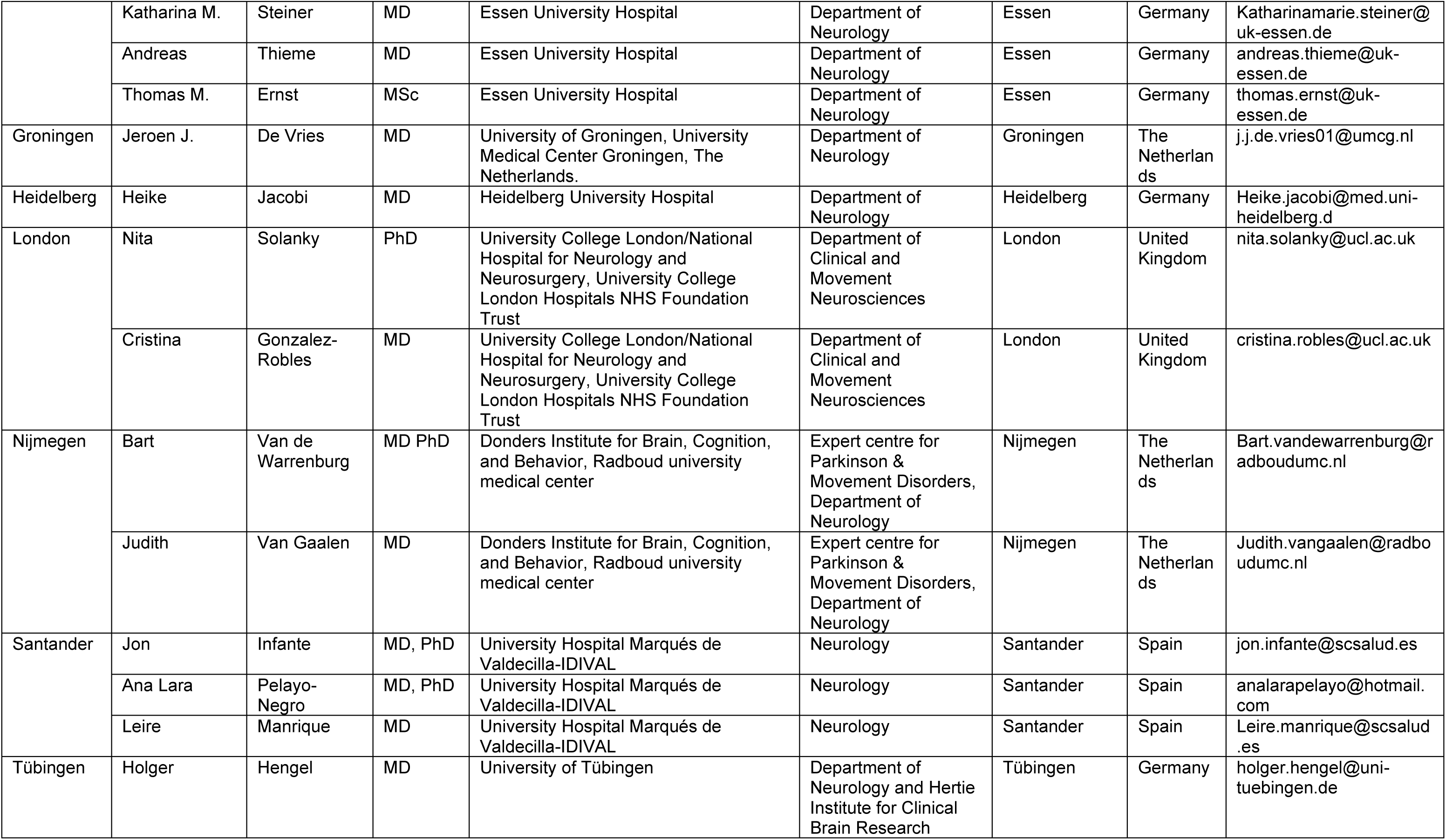
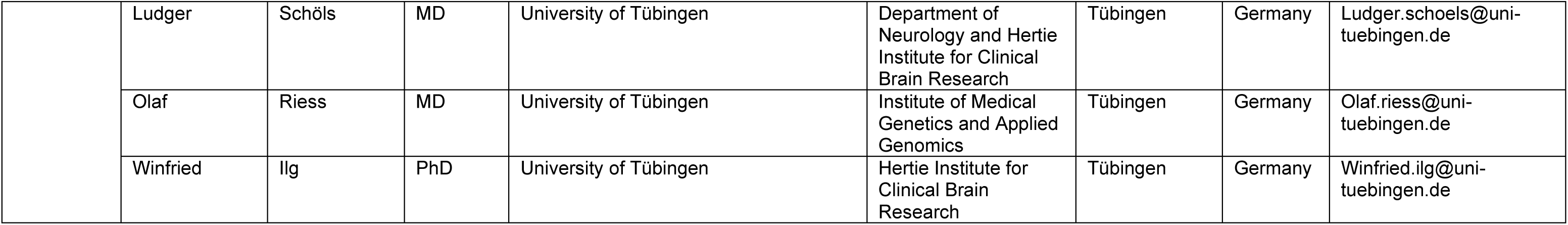

## Supplementary Figures

**Supplementary Figure 1.**
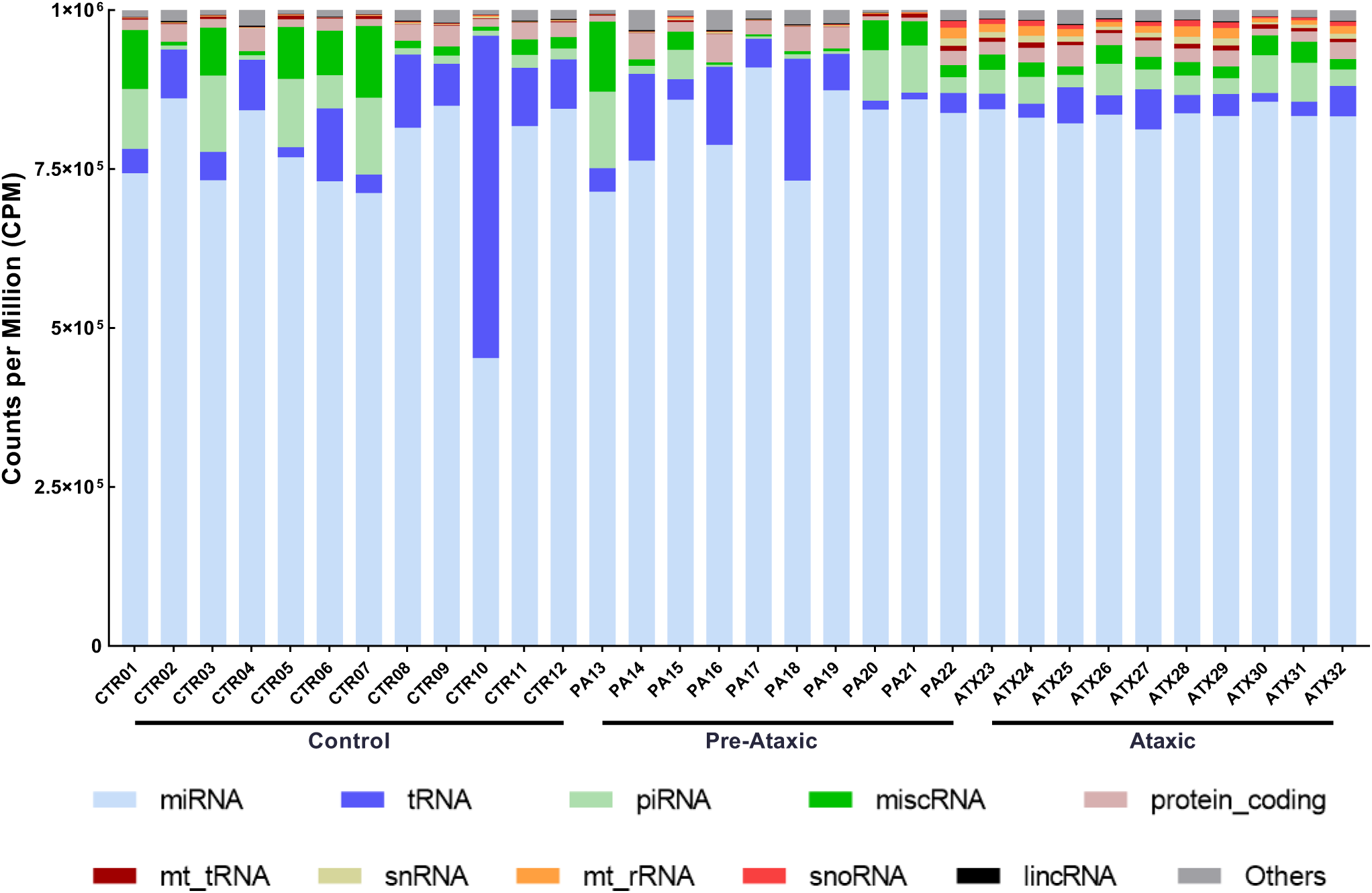
Distribution of RNA biotypes in EVs from each participant of the study. This figure depicts the proportion of mapped RNA classes in each library. Control #10 was excluded from further analysis due to the distinct RNA composition.

**Supplementary Figure 2.**
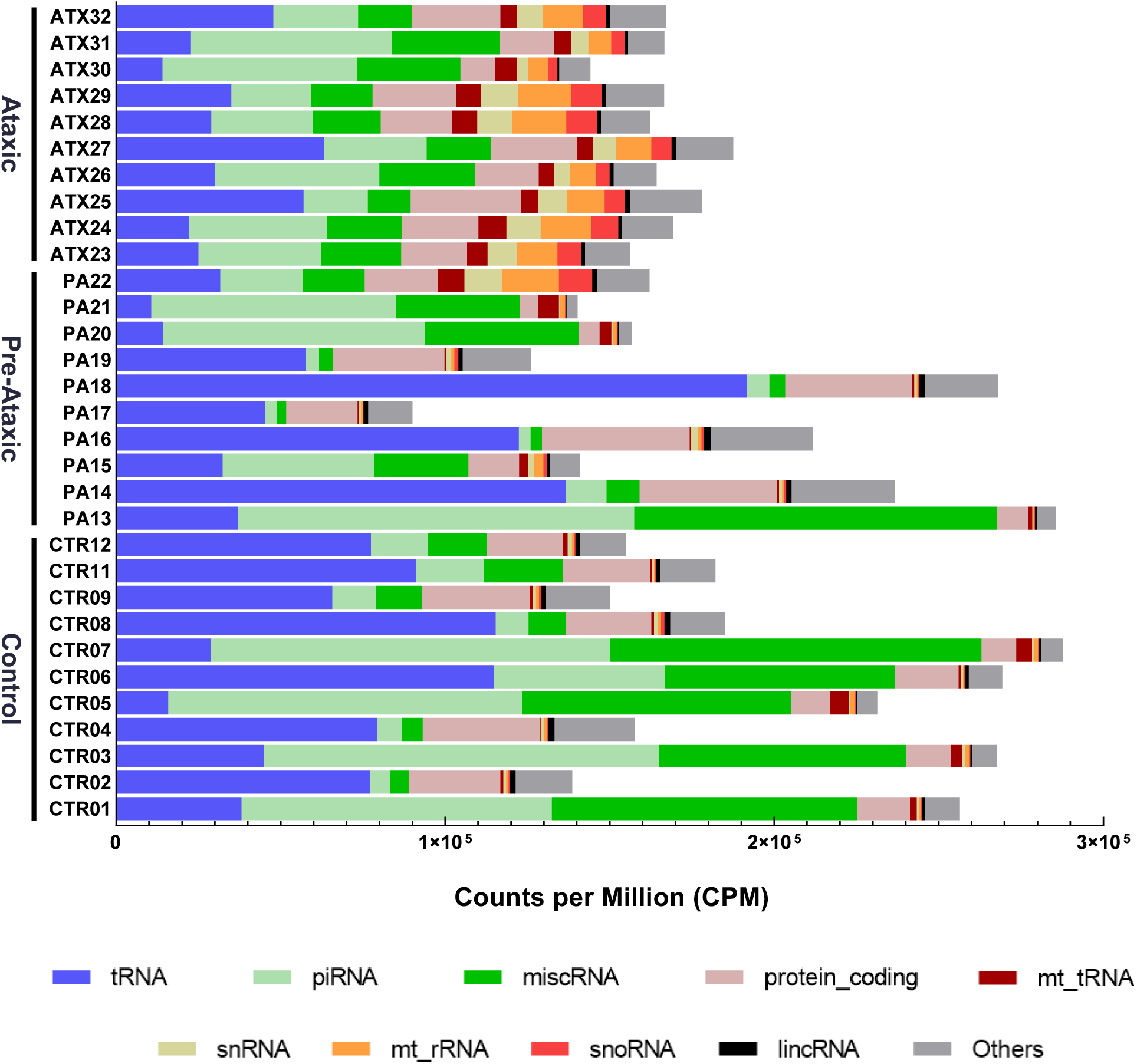
RNA biotypes distribution in EVs (excluding miRNAs). This figure illustrates the distribution of RNA biotypes within extracellular vesicles (EVs), specifically excluding miRNA content. Each bar chart represents the relative proportion of mapped RNA classes in each library, providing a comprehensive view of RNA diversity within EVs, while omitting miRNA data for focused analysis.

**Supplementary Figure 3.**
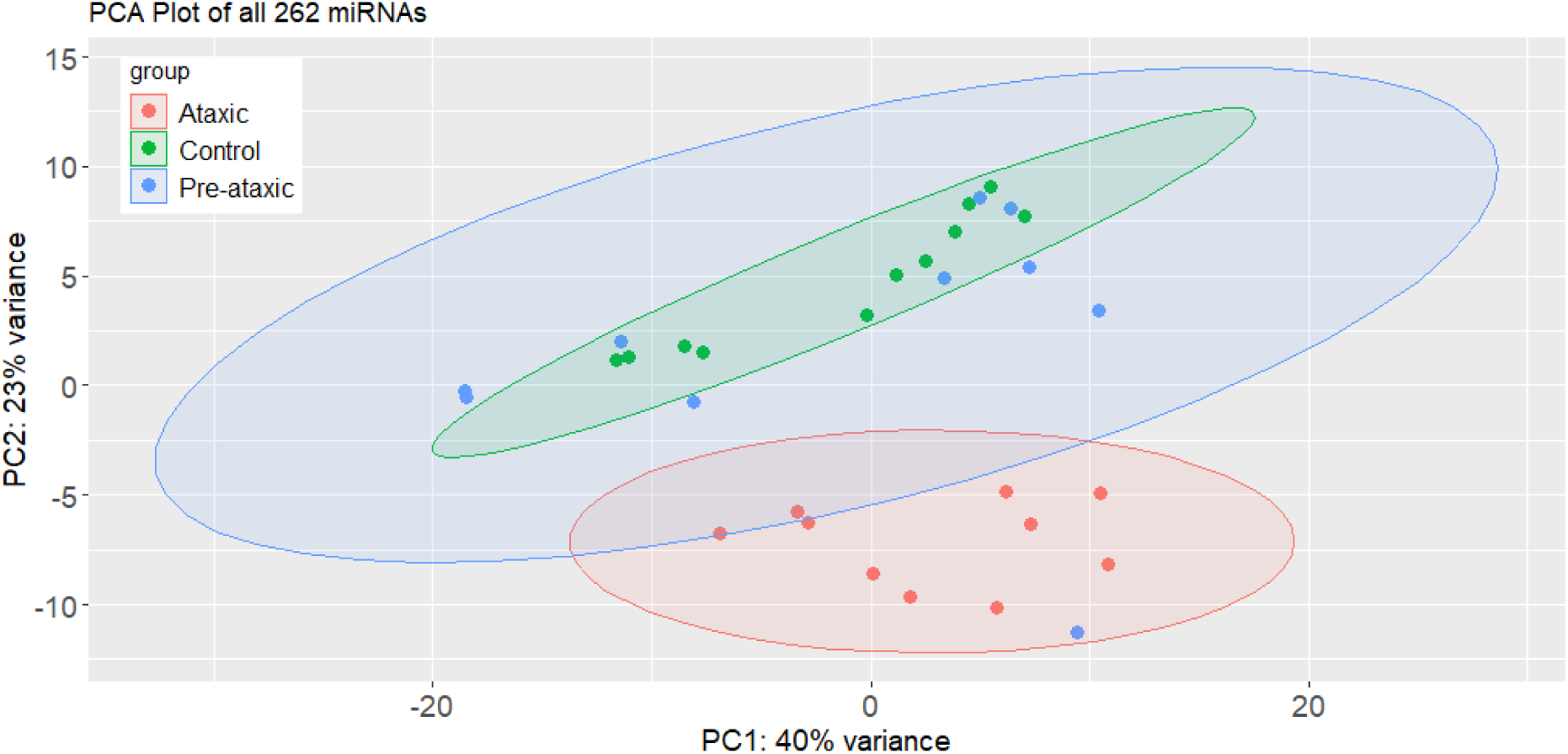
PCA plot of 262 EVs-miRNA in controls, pre-ataxic and ataxic SCA3 mutation carriers. Each point represents an individual sample colored by their respective group. Ataxic group is highlighted in red. The plot shows a clear separation between ataxic and the remaining study population.

**Supplementary Figure 4.**
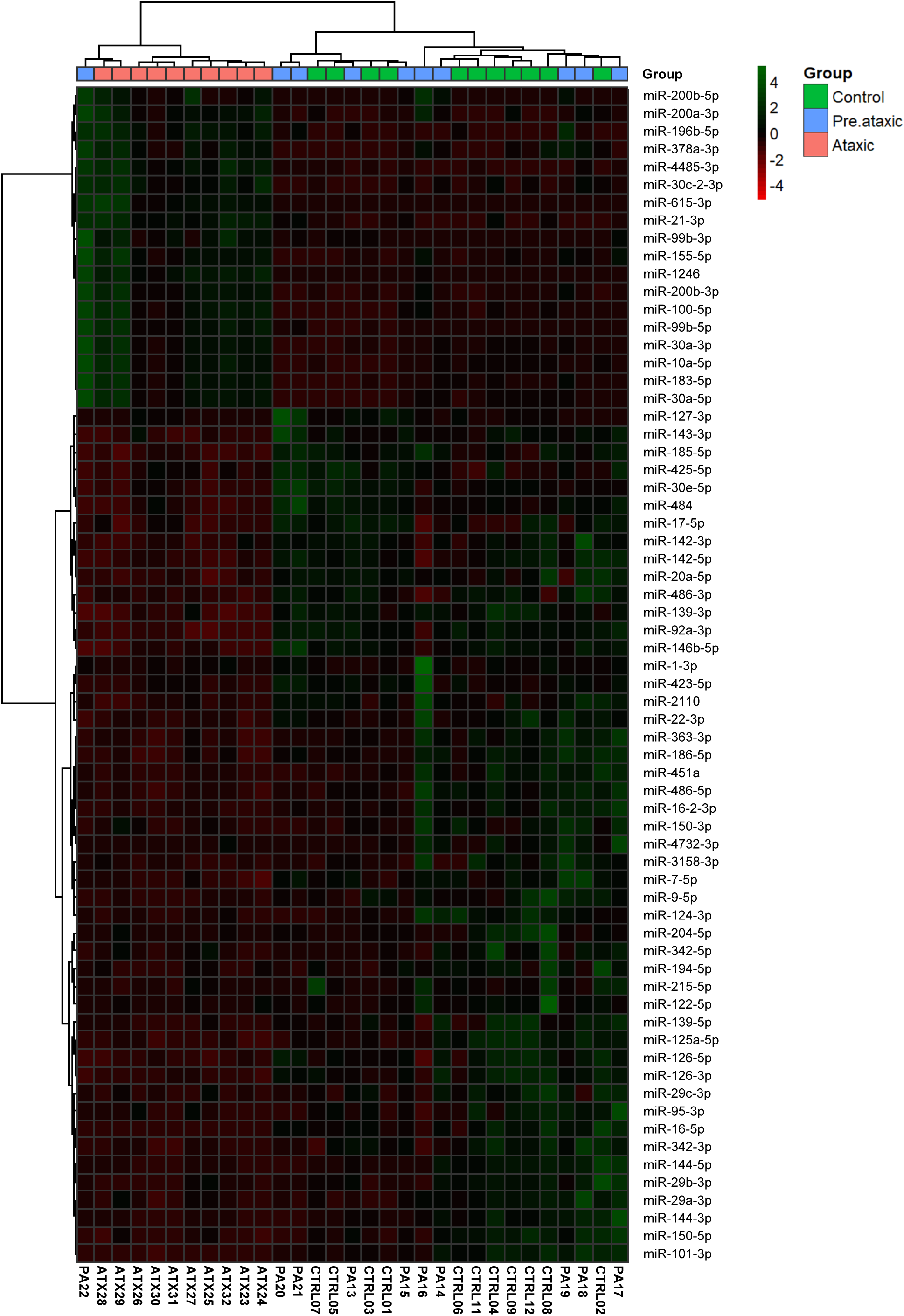
Hierarchical cluster analysis of DE miRNAs. The heatmap displays the clustering of the 66 significant DE miRNAs identified in the individual pairwise comparison between ataxic mutation carriers and controls (53), and between pre-ataxic carriers and ataxic participants (13 that did not overlap with the previous 53 miRNAs; see Figure 3D). The analysis used z-scores of the expression levels of each individual for each miRNA. The dendrogram illustrates the clustering of the different individuals and miRNAs based on Euclidean distance as the measure of dissimilarity, using Ward’s minimum variance linkage method.

**Supplementary Figure 5.**
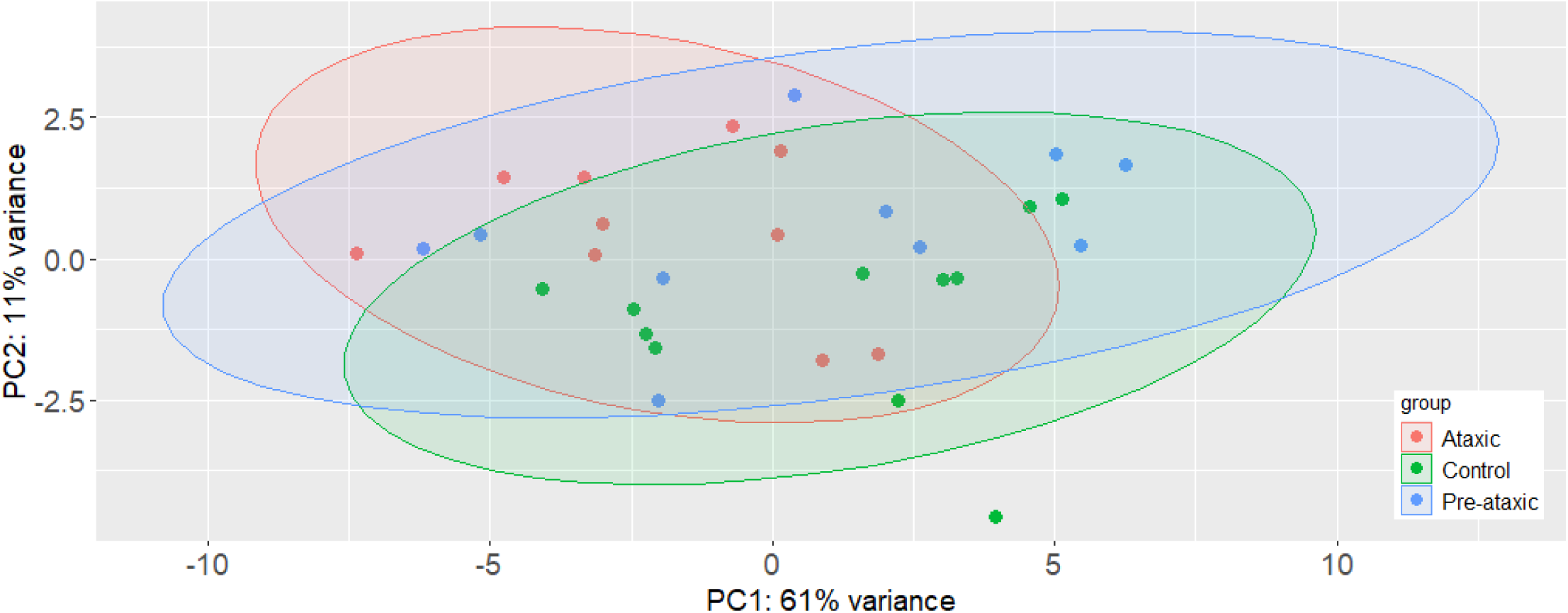
PCA plot of 22 tRNAs contained in EVs of controls, pre-ataxic and ataxic SCA3 mutation carriers. Each data point represents an individual sample and is colored according to its corresponding group. Ataxic group is highlighted in red, pre-ataxic in blue and controls in green.

**Supplementary Figure 6.**
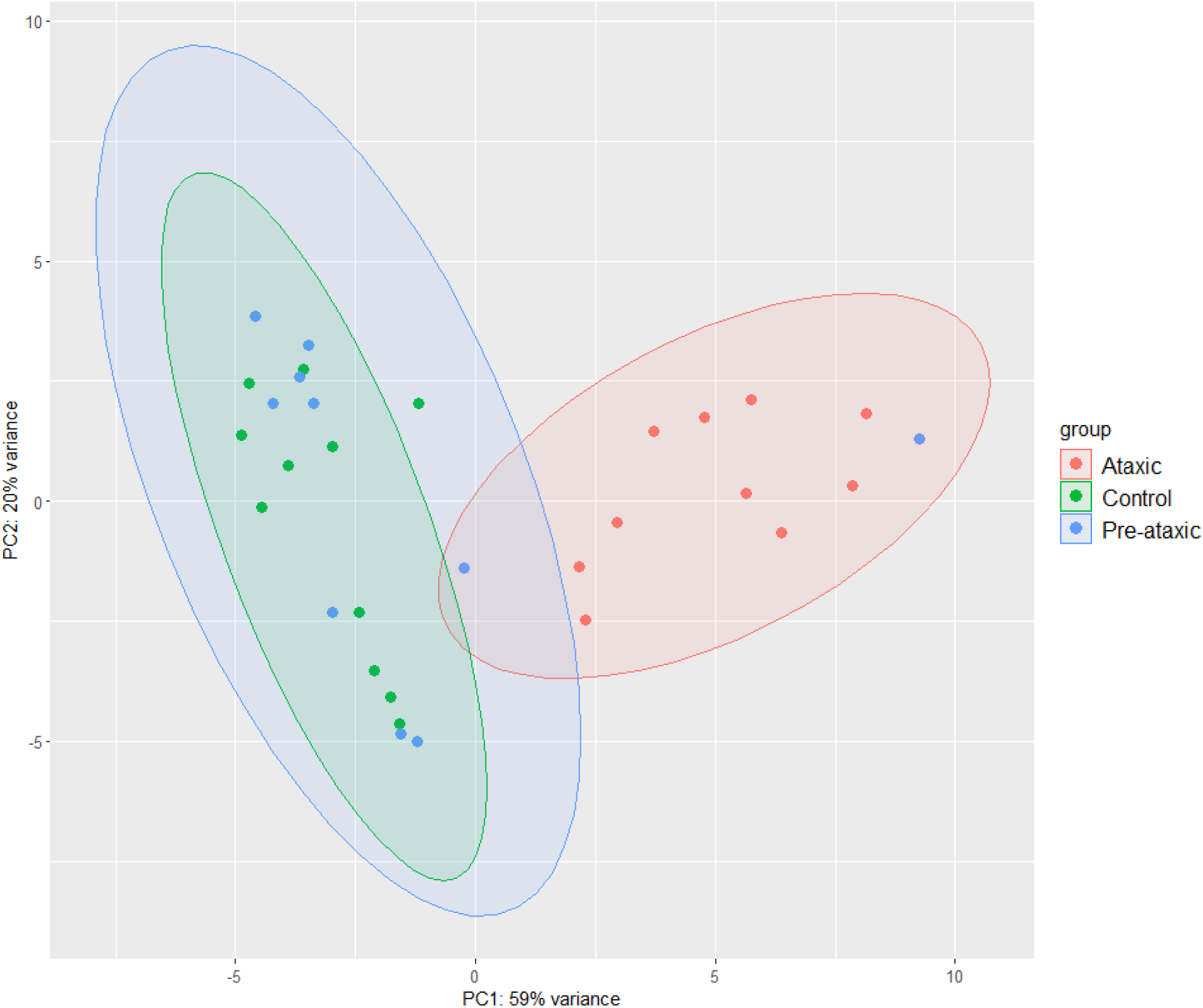
PCA plot of 20 piRNAs contained in EVs of controls, pre-ataxic and ataxic SCA3 mutation carriers. Each data point represents an individual sample and is colored according to its corresponding group. Ataxic group is highlighted in red, pre-ataxic in blue and controls in green. piRNAs plot demonstrate a distinct separation between ataxic samples and the rest of the study population.

